# Conserved *cis* and *trans* communication domains mediate module interaction in glycopeptide antibiotic NRPS assembly lines

**DOI:** 10.64898/2026.08.07.743486

**Authors:** Thomas M. Hamm, Dardan Beqaj, Nina Pfahler, Andreas Kulik, Thilo Stehle, Wolfgang Wohlleben, Evi Stegmann

**Affiliations:** Microbial Active Compounds, Interfaculty Institute of Microbiology and Infection Medicine (IMIT), University of Tübingen, Auf der Morgenstelle 28, 72076 Tübingen, Germany; Interfaculty Institute of Biochemistry, University of Tübingen, Auf der Morgenstelle 34, 72076 Tübingen, Germany; Microbiology/Biotechnology, Interfaculty Institute of Microbiology and Infection Medicine (IMIT), University of Tübingen, Auf der Morgenstelle 28, 72076 Tübingen, Germany; Partner-Site: DZIF Tübingen, Auf der Morgenstelle 28, 72076 Tübingen, Germany; Excellence Cluster "Controlling Microbes to Fight Infections" (CMFI), University of Tübingen, 72076 Tübingen, Germany

**Keywords:** Natural products, non-ribosomal peptide synthetase, actinomycetes, antibiotics, glycopeptides, COM domains, docking domains

## Abstract

Non-ribosomal peptide synthetases (NRPSs) assemble structurally complex and clinically important natural products, yet the mechanisms coordinating communication between multiple enzymes remain incompletely understood. Here, we dissected the NRPS system underlying biosynthesis of the glycopeptide antibiotic (GPA) balhimycin in *Amycolatopsis balhimycina* to elucidate principles of inter-enzyme communication. Genetic perturbation of short terminal structural elements markedly reduced balhimycin production, demonstrating their critical role in biosynthesis. AlphaFold3 predictions identified these elements as *trans*-COM domains mediating specific NRPS interactions, primarily through hydrophobic contacts. Quantitative binding studies using microscale thermophoresis confirmed the importance of the *trans*-COM domains for multiprotein interaction, extending current models of NRPS communication. Comparative structural analysis further uncovered a conserved class of *cis*-COM domains within condensation domains across GPA NRPS assembly lines. Together, our findings establish a unified model in which *trans*- and *cis*-interfaces cooperatively maintain assembly line fidelity, redefining NRPS architecture and enabling rational engineering strategies.

## Introduction

Non-ribosomal peptide synthetases (NRPSs) produce a wide range of clinically important natural products, including glycopeptide antibiotics (GPAs) like vancomycin and teicoplanin^1^. These are classified into five structural types (Types I-V) according to their backbone composition, characteristic crosslinking patterns and tailoring modifications^1^. The pathway leading to the production of the vancomycin-type I antibiotic balhimycin in *Amycolatopsis balhimycina* has been extensively characterized, as it was among the first genetically tractable GPA systems^2–4^. In this pathway, three NRPSs (BpsA-C) assemble a heptapeptide backbone composed of both proteinogenic and non-proteinogenic amino acids in a strictly defined sequence^4–6^ (Figure 1). The fidelity of this process critically depends on precise and selective interactions between individual NRPS subunits, as errors in subunit coordination would compromise product formation.

**Figure 1.**
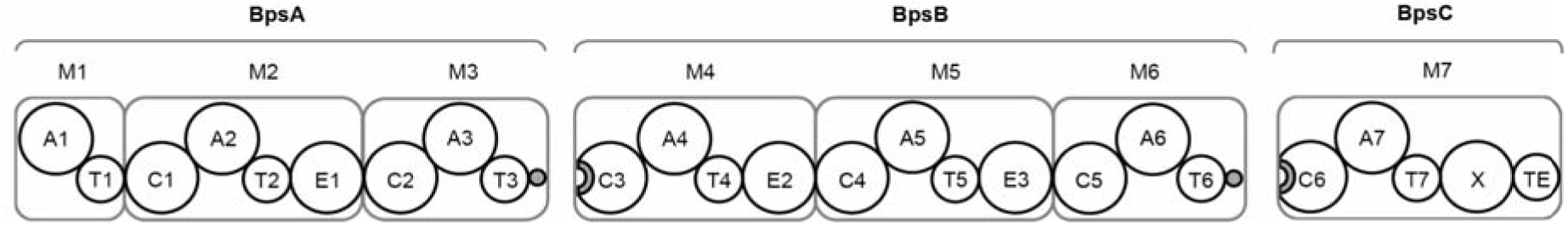
NRPS assembly line of balhimycin biosynthesis in *A. balhimycina*. Schematic representation of the modular balhimycin NRPSs BpsA-C. Grey dots and semicircles indicate the putative interacting regions. A, adenylation domain; T, thiolation domain; C, condensation domain; E, epimerization domain; X, X domain; TE, thioesterase domain; M, module. Numbers indicate the domain order.

Such coordination represents a general challenge in NRPS assembly lines, which operate in a modular, assembly-line fashion. Each module minimally comprises adenylation (A), thiolation (T), and condensation (C) domains responsible for substrate activation, carrier tethering, and peptide bond formation while optional epimerization (E) domains generate D-amino acid residues^7–10^. In GPA biosynthesis, specialized X domains located in the terminal modules recruit the cytochrome P450 monooxygenases responsible for oxidative crosslinking reactions that generate the characteristic three-dimensional GPA scaffold^11–14^. A terminal thioesterase (TE) domain catalyzes the release of the mature peptide from the NRPS assembly line.

The directional transfer of intermediates between modules encoded on separate proteins requires highly specific protein-protein interactions to ensure both efficiency and fidelity while preventing non-productive crosstalk.

Bioinformatic analysis of the MIBiG repository indicated that GPA NRPS assembly lines contain communication-mediating (COM) domains^15^. These domains occur in two organizational forms: *trans*-COM and *cis*-COM domains^16,17^. In *trans*-COM domains, the interaction interface is split between two sequentially acting NRPS proteins. A donor region (COM^D^) is located at the C terminus of an upstream NRPS, whereas an acceptor region (COM^A^) is embedded within the N-terminal C domain of the downstream NRPS. Binding between COM^D^ and COM^A^ promotes association of the two NRPS proteins, thereby facilitating communication between adjacent modules.

Recently, the first structure of a *trans*-COM domain complex was reported^16^, confirming a helix-hand interaction model in which COM^D^ forms an α-helix that binds to a β-hand-like COM^A^ element within the downstream C domain. Notably, the structure also revealed direct contacts between the *trans*-COM domain and the acceptor T domain, involving residues from both COM^D^ and COM^A^. These findings suggest that *trans*-COM domains not only mediate association between NRPS proteins but may also contribute to recruitment and positioning of the downstream T domain. *Trans*-COM domains have been identified in two different NRPS domain architectures, termed E-COM^D^/C and T-COM^D^/C^15–18^. In these arrangements, COM^D^ is directly C-terminal to either an E domain or a T domain and mediates interaction with the C domain of the downstream NRPS protein.

In contrast, *cis*-COM domains contain the helix and hand elements within a single continuous polypeptide. To date, *cis*-COM domains have been observed only in E-COM-C domain topologies. Both *cis*- and *trans*-COM domains are proposed to serve a scaffolding function between E and C domains in NRPS assembly lines^17^.

Although COM domains are known to play important roles in organizing NRPS assembly lines, their functional and structural contributions to GPA biosynthesis remain poorly understood.

In the balhimycin system, the NRPS subunits BpsA-C must interact with high specificity to assemble the heptapeptide scaffold, yet the molecular basis of these interactions has remained unclear.

Here, we combined genetic, biochemical, and computational approaches to dissect inter-enzyme communication in the balhimycin NRPS system and to define the broader role of COM domains in GPA biosynthesis. Our results reveal the structural basis of NRPS subunit interactions in the balhimycin assembly line and identify COM domains as conserved intermodular interfaces in GPA biosynthetic systems. Notably, we identify a previously unknown COM domain topology. Together, these findings provide mechanistic insight into the organization of GPA assembly lines and establish a foundation for the rational engineering of GPA biosynthesis.

## Results

### C-terminal truncation of BpsA and BpsB reduces but does not abolish balhimycin production

To elucidate the functional role of the C-terminal structural elements putatively mediating NRPS-NRPS interaction (BpsA-BpsB; BpsB-BpsC) during balhimycin biosynthesis, we generated targeted truncation mutants in *A. balhimycina*, namely *A. balhimycina* DB1, lacking the C-terminal 37 amino acids of BpsA; *A. balhimycina* DB2, lacking the C-terminal 36 amino acids of BpsB; and *A. balhimycina* DB1-2, a double mutant lacking both corresponding C-terminal regions.

Culture filtrates from the three mutants exhibited significantly reduced inhibitory activity against *Bacillus subtilis* 168 (Figure 2b). Inhibition zones were detectable only after 120 hours of cultivation, that is, considerably later than observed for the wild-type culture filtrate, which displayed an inhibition zone already after 48 hours. To confirm that this inhibition was due to balhimycin, we performed high-performance liquid chromatography (HPLC) coupled with electrospray ionization mass spectrometry (HPLC-ESI-MS) (Figure 2a). In culture filtrates of all three mutants, peaks corresponding to balhimycin appeared at retention times of 5-5.5 minutes, consistent with the wildtype profile (Figure S1). We identified peaks with *m/z* 1305.34 [M+H]^+^ for mono-glycosylated and *m/z* 1446.46 [M+H]^+^ for bi-glycosylated balhimycin (Figure S1). Notably, in the three mutants, mono-glycosylated balhimycin (*m/z* 1305.34 [M+H]^+^) predominated, except in *A. balhimycina* DB1, which, in addition, exhibited a relatively higher proportion of bi-glycosylated balhimycin (Figure S1). Quantitative analysis of the HPLC-ESI-MS data revealed a positive correlation between antibiotic activity and balhimycin production levels (Figure 2a,b), *A. balhimycina* DB1 produced slightly higher amounts than *A. balhimycina* DB2 and *A. balhimycina* DB1-2, but all mutants exhibited significantly reduced production compared to the wild type.

**Figure 2.**
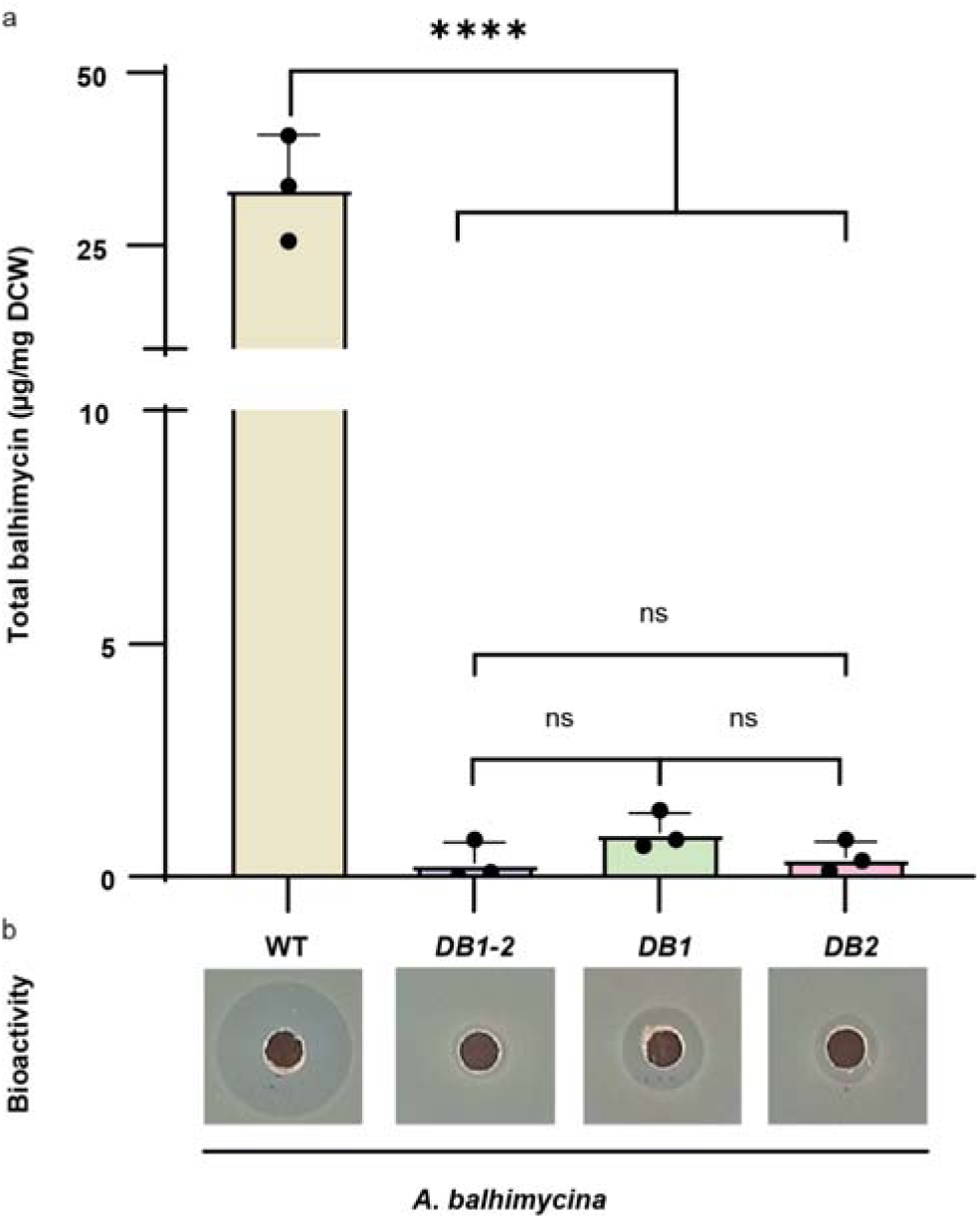
HPLC-MS-based quantification and bioactivity test of balhimycin production. (**a**) Quantified amounts of balhimycin produced by the mutants *A. balhimycina DB1*, lacking the C-terminal 37 amino acids of BpsA; *A. balhimycina DB2*, lacking the C-terminal 36 amino acids of BpsB; and *A. balhimycina DB1-2* lacking both regions. Error bars indicate standard deviations of three biological replicates, (****) p<0.05; (ns) not significant. DCW: Dry cell weight; ns: not significant.(**b**) Bioactivity assay against *B. subtilis 168* exposed to the culture filtrate of the wild-type (WT) and respective mutants.

These results indicate that the C-terminal regions of BpsA and BpsB are not essential for balhimycin biosynthesis but are crucial for efficient production.

### Structural modeling revealed a high-confidence α-helix/linker-mediated interface between adjacent balhimycin NRPS modules

To investigate the molecular basis underlying the reduction in balhimycin biosynthesis in the three mutants compared to the wildtype, we used AlphaFold 3 (AF3) to model the interfaces between the C-terminal regions of BpsA and BpsB and the N-terminal regions of the downstream NRPS proteins, BpsB and BpsC, respectively. For the BpsA-BpsB interaction, we analyzed the C-terminal 117 amino acids of BpsA, including the terminal T3 domain, together with the N-terminal 436 amino acids of BpsB containing the C3 domain (Figure 1). For the BpsB-BpsC interaction, we modeled the C-terminal 113 amino acids of BpsB, comprising the T6 domain, together with the N-terminal 465 amino acids of BpsC including the C6 domain.

In both cases, AF3 predicted that each C-terminal region contains two structural elements, a T domain and a C-terminal α-helix connected via an unstructured linker, which contact the downstream C domain (Figure 3a, b). Across 25 models from five independent runs, the α-helix and part of the linker were consistently placed at the same site on the C domain, whereas the T domain adopted multiple positions (Figure 3c, d). This variability was also reflected in the corresponding predicted aligned error (PAE) plots (Figure 4a, b). For the BpsA-BpsB interface, four alternative positions were observed for the T3 domain, with the highest-confidence models placing the T3 domain in one predominant position (Figure 3c, 1).

**Figure 3.**
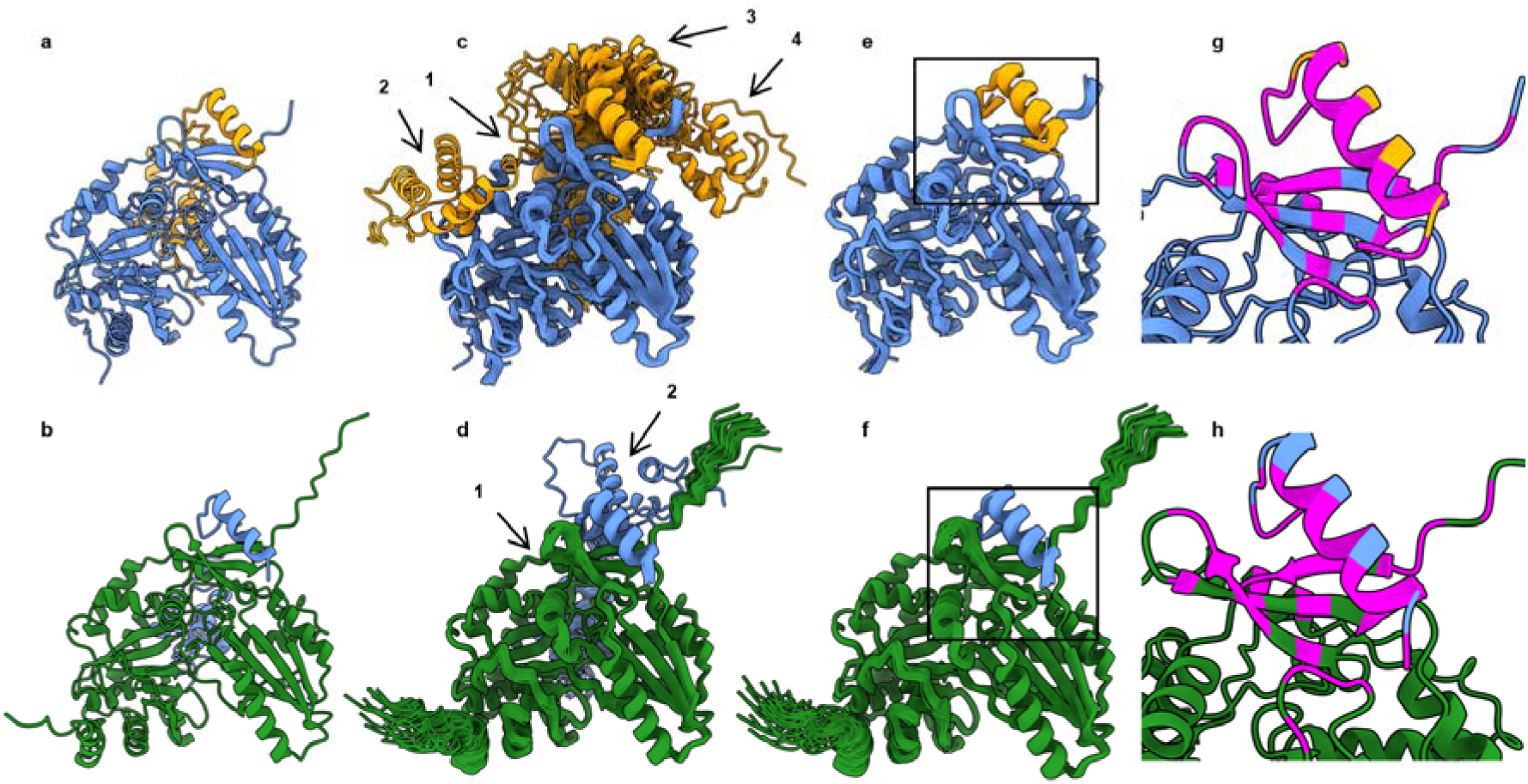
Predicted interactions between the C-terminal regions of BpsA (orange) and BpsB (blue) with the N-terminal regions of BpsB and BpsC (green). (**a**, **c**) Predicted interactions between the C-terminal region of BpsA (orange; T domain - linker - helix) and the N-terminal C domain of BpsB (blue). (**a**) shows one AF3 predicted model, while (**c**) displays 25 superimposed predicted models. The C-terminal α-helix of BpsA is consistently predicted to bind at the same site of the N-terminal C domain of BpsB. In contrast, the C-terminal T domain of BpsA is predicted to interact with four distinct sites on the N-terminal C domain of BpsB (1-4). (**b**, **d**) Predicted interactions between the C-terminal part of BpsB (blue; T domain - linker - helix) and the N-terminal C domain of BpsC (green). The one model (**b**) as well as the 25 superimposed models (**d**) show similar results as before, except that the C-terminal T domain of BpsB is predicted to bind at only two distinct sites on the N-terminal C domain of BpsC (1, 2). The 25 superimposed models of predicted interactions between the C-terminal helix and linker of BpsA and the N-terminal C domain of BpsB (**e**) show very similar relative orientations of the two structural elements. The same can be seen in the depiction of the 25 superimposed models of predicted interactions between the C-terminal helix and linker of BpsB and the N-terminal C domain of BpsC (**f**). (**g**, **h**) Visualization of the interface analysis results from PISA. The amino acids of the binding interfaces of the C-terminal helix of BpsA (orange) with the N-terminal C domain of BpsB (blue) (**g**) and the C-terminal helix of BpsB with the N-terminal C domain of BpsC (green) (**h**) are colored in magenta. All the C domains of BpsB and BpsC are depicted in the same orientation (**a**-**f**). The rectangle (**e**, **f**) shows the magnified area in (**g**, **h**), respectively.

**Figure 4.**
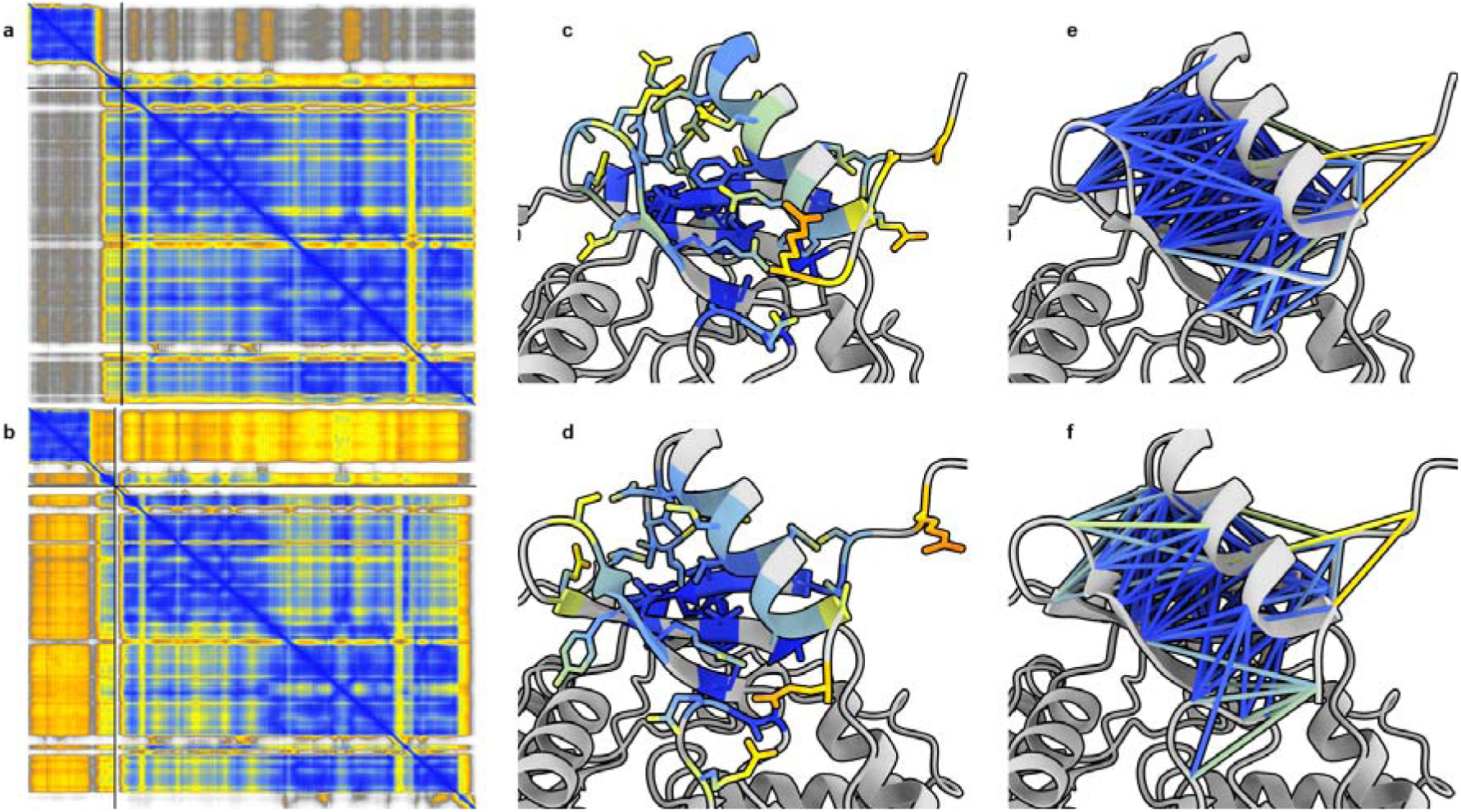
PAE and pLDDT of predicted structures of the C-terminal parts of BpsA and BpsB with the N-terminal parts of BpsB and BpsC. The PAE plots (**a**, **b**) belong to the structures depicted in Figure 3a and b. The plots are representative of the 50 PAE plots of all the 25 predicted interactions between the C-terminus of BpsA with the N-terminus of BpsB and the C-terminal part of BpsB with the N-terminal part of BpsC (Figure 3c, **d**). (The PAE is visualized according to the following color scheme: dark blue 0 Å, light blue 5 Å, yellow 10 Å, orange 15 Å, dark gray 20 Å, gray 25 Å, and white 30 Å.) In **c** and **d** the predicted confidence of the atoms in the binding interface between the C-terminal helix of BpsA with the N-terminal C domain of BpsB (**c**) and the C-terminal helix of BpsB and the N-terminal C domain of BpsC (**d**) is depicted via the pLDDT. (In **c** and **d** only the side chains of amino acids involved in the interface are shown and the backbone depicted as cartoon is colored in grey. The pLDDT is visualized according to the following color scheme: dark blue 100, light blue 90, yellow 70, orange 50, and red 0.) In **e** and **f** the amino acids of the interface between the C-terminus of BpsA and the N-terminal C domain of BpsB (**e**) and the C-terminus of BpsB and the N-terminal C domain of BpsC (**f**) with a distance of up to 8 Å to each other are connected via a line. The colors of these lines show the PAE of the respective residue pair.

For the BpsB-BpsC interface, the T6 domain was predicted mainly in one position but with overall lower interface confidence (Figure 3d). Thus, the helix/linker interaction was robustly reproduced, whereas the relative arrangement of the T and C domains remained uncertain.

To obtain high-quality models suitable for detailed subsequent structural analysis, we examined the two C-terminal elements separately. AF3 predictions of the C-terminal α-helix/linker segments of BpsA (21 amino acids) or BpsB (20 amino acids) with the corresponding C domains yielded highly consistent models across all 25 predictions for each interface (Figure 3e, f). These consistently exhibited high scores for the predicted template modelling (pTM > 0.78) and interface pTM (ipTM > 0.87), indicating high-confidence binding modes. In contrast, isolated T domains produced multiple alternative poses with low and variable interface confidence (Figure S2), largely inconsistent with those observed in the full-length models (Figure 3c, d). Together, these results identify the α-helix/linker-mediated contacts at the BpsA-BpsB and BpsB-BpsC interfaces as the only interaction modes robustly supported by AF3 with high confidence, providing a reliable basis for further structural analysis.

### Residue-level analysis supports a conserved *trans*-COM domain interface between adjacent balhimycin NRPS proteins

To characterize the α-helix/linker-mediated interfaces in more detail, we analyzed the top-ranked AF3 models using PISA and AF3 confidence metrics. For the BpsA-BpsB interface, PISA analysis of the top-ranked structures from five independent AF3 runs showed only minor variation in the interacting residues. Of the 21 C-terminal residues of BpsA, 15-16 consistently contributed to the interface. In the N-terminal C3 domain of BpsB 24-26 residues were engaged, primarily located within a β-sheet and adjacent loops, with additional contributions from a neighboring helix and loop region (Table S1, Figure 3g). The corresponding predicted local distance difference test (pLDDT) values were predominantly high within the interface core (Figure 4c), while the PAE values indicated low predicted relative positional errors for most contacting residues (Figure 4e).

A similar pattern was observed for the BpsB-BpsC interface. Across the top-ranked models from five independent AF3 runs, 12-14 of the 20 C-terminal residues of BpsB and the same set of 24 residues in the N-terminal C6 domain of BpsC consistently formed the interface (Table S1, Figure 3h). As in BpsA-BpsB, the interacting residues in the C6 domain of BpsC were located primarily within a β-sheet and adjacent loops, with additional contributions from a nearby helix and loop region. Interface residues again showed predominantly high pLDDT values (Figure 4d) and low PAE values, consistent with a reliable residue-level prediction (Figure 4f).

Together, these analyses show that the BpsA-BpsB and BpsB-BpsC interfaces share the same overall architecture: a C-terminal α-helix and part of a linker bound to a β-sheet-containing surface of the downstream N-terminal C domain. This structural motif corresponds to the *trans*-COM domain architecture described in other NRPS systems^16^. The high reproducibility of the interacting residues across independent AF3 runs, together with favorable confidence metrics, supported the robustness of the predicted interfaces for further structural analysis.

### Hydrophobic and electrostatic complementarity support partner-adapted interfaces

To further assess the plausibility of the AF3-predicted interfaces, we analyzed hydrophobic and electrostatic surface complementarity together with PISA-derived hydrogen bonds, salt bridges and interface areas. Hydrophobic surface mapping showed that the COM^A^ regions within the N-terminal C domains of both BpsB and BpsC present two main hydrophobic features at the interface: a small, conserved pocket and a larger adjacent groove formed primarily by β-sheet residues and neighboring structural elements (Figure 5c, d). The pocket is formed by the same amino acids in both proteins, whereas the larger hydrophobic groove differs subtly between the two COM^A^ regions, including a broader, more contoured surface in BpsB and a narrower groove in BpsC.

**Figure 5.**
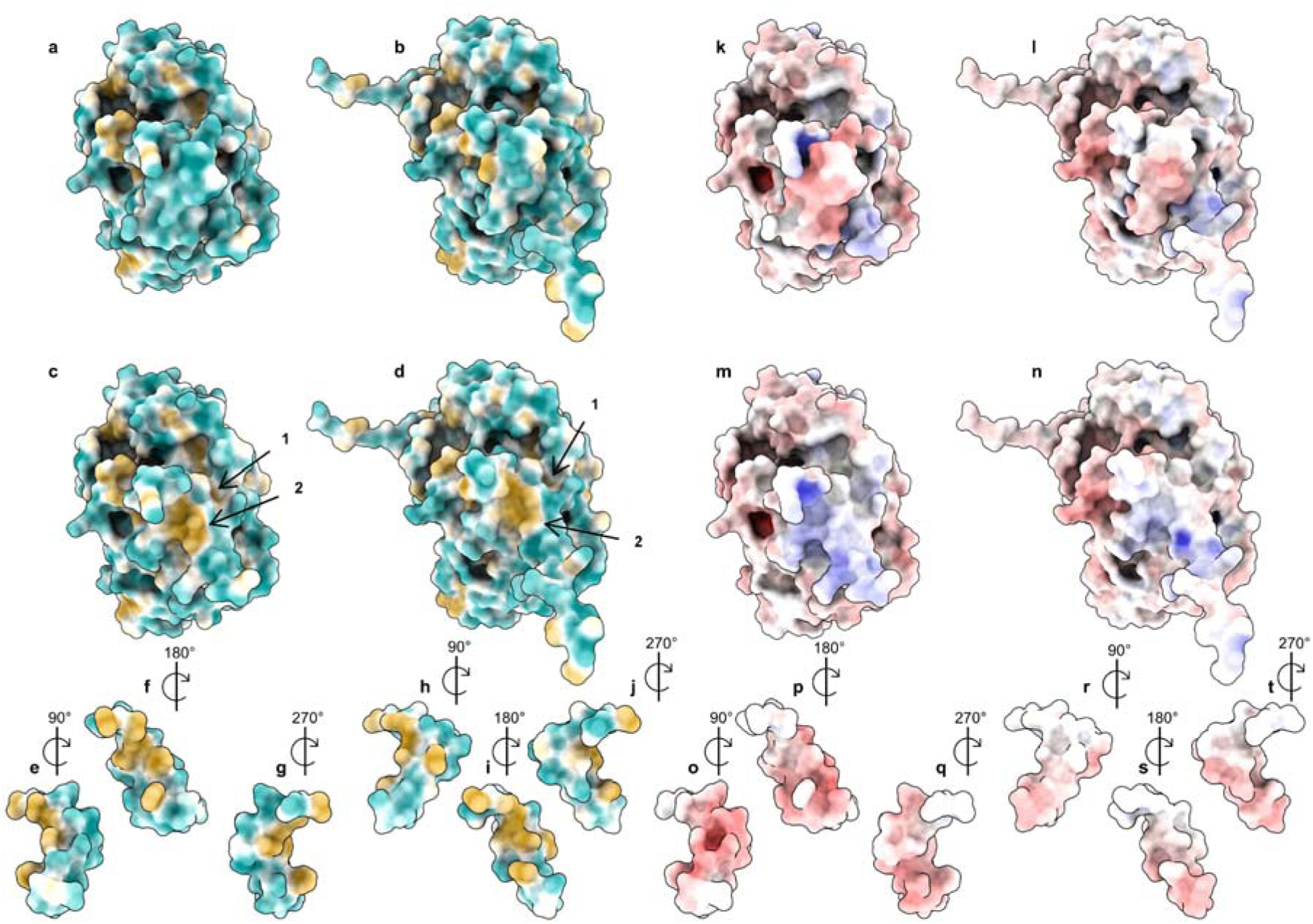
Visualization of the molecular lipophilicity potential and the electrostatic potential on the solvent-excluded surfaces of the two predicted *trans*-COM domains. The surfaces of the COM^D^ region of BpsA bound to the COM^A^ region within the N-terminal C domain of BpsB (**a**, **c**) and the C-terminal COM^D^ region of BpsB bound to the COM^A^ region within the N-terminal C domain of BpsC (**b**, **d**) colored by the molecular lipophilicity potential. The COM^D^ region is shown in (**a**) and (**b**), but not in (**c**) and (**d**), to show the COM^A^ region binding sites. The coloring ranges from dark cyan for the hydrophilic parts, through white, to dark goldenrod for the lipophilic parts. The arrows point at the small conserved hydrophobic pocket (1) and the larger adjacent hydrophobic groove (2). (**e**-**j**) The structures of the COM^D^ regions are taken from the predicted protein-protein interactions shown in (**a** and **b**) and are turned right by 90° (**e**, **h**), 180° (**f**, **i**) and 270° (**g**, **j**) to show the residues involved in the protein-protein interaction. The surfaces of the COM^D^ region of BpsA bound to the COM^A^ region within the N-terminal C domain of BpsB (**k**, **m**) and the C-terminal COM^D^ region of BpsB bound to the COM^A^ region within the N-terminal C domain of BpsC (**l**, **n**) colored by electrostatic potential. The COM^D^ region is shown In (**k**) and (**l**), but not in (**m**) and (**n**), to show the COM^A^ region binding sites. A positive electrostatic potential is shown in blue and a negative one in red. (**o**-**t**) The structures of the COM^D^ regions are taken as in the predicted protein-protein interactions shown in (**k** and **l**) and are turned right by 90° (**o**, **r**), 180° (**p**, **s**) and 270° (**q**, **t**) to show the residues involved in the protein-protein interaction.

In both interfaces, hydrophobic residues from the COM^D^ region dock into these conserved surface features. In particular, three hydrophobic linker residues contributed similarly in BpsA and BpsB, indicating a conserved mode of linker-mediated packing (Figure 5e-j). By contrast, the α-helical contributions differed between the two interfaces and matched the distinct surface topology of the downstream COM^A^ regions. Thus, although both interfaces share the same overall hydrophobic interaction principle, they appear to be locally adapted to the specific shape of the respective COM^A^ regions surface.

Electrostatic surface calculations revealed complementary charge distributions at both interfaces (Figure 5k-t). The surfaces of the COM^A^ regions were predominantly positively charged (Figure 5m, n), whereas the interacting COM^D^ regions of BpsA and BpsB were largely negatively charged (Figure 5o-t), consistent with electrostatic attraction. This complementarity was more pronounced in the BpsA-BpsB interface, which showed a stronger positive potential on BpsB and a stronger negative potential on BpsA, whereas the BpsB-BpsC interface displayed a weaker and more spatially restricted charge contrast.

PISA analysis further supported both interfaces quantitatively. For the BpsA-BpsB *trans*-COM domain, the top-ranked models from five AF3 runs contained on average 11.4 hydrogen bonds and 10.2 salt bridges, with an average interface area of 866.0 Å^2^, corresponding to 37.0% of the solvent-accessible surface of the BpsA COM^D^ region (Table S1). For the BpsB-BpsC *trans*-COM domain, the corresponding values were 8.2 hydrogen bonds, 4.6 salt bridges and 755.7 Å^2^, corresponding to 35.8% of the solvent-accessible surface of the BpsB COM^D^ region.

Together, these analyses provide a plausible structural explanation for stabilization of the predicted interfaces through hydrophobic packing and electrostatic complementarity. Although the interfaces are broadly similar, their local differences suggest a degree of partner adaptation that may contribute to interaction specificity.

### COM^D^ regions mediate selective intermodular recognition

To further investigate the role of the *trans*-COM domains in intermodular recognition, we generated six protein constructs and analyzed their interactions *in vitro* by microscale thermophoresis (MST). T3-COM^D^_BpsA_ and T6-COM^D^_BpsB_ comprised the terminal T domain and the C-terminal COM^D^ region of BpsA and BpsB, respectively. C3_BpsB_ and C6_BpsC_ comprised the N-terminus through the end of the first C domain of BpsB and BpsC, including the COM^A^ regions. T3-ΔCOM^D^_BpsA_ and T6-ΔCOM^D^_BpsB_ contained the terminal T domains of BpsA and BpsB but lacked the C-terminal COM^D^ regions.

The cognate donor-acceptor combinations T3-COM^D^_BpsA_/C3_BpsB_ and T6-COM^D^_BpsB_/C6_BpsC_ yielded K_D_^app^ values of approximately 250 μM and 320 μM, (Figure 6a, c). The corresponding non-cognate combinations, T3-COM^D^_BpsA_/C6_BpsC_ and T6-COM^D^_BpsB_/C3_BpsB_, showed substantially weaker interaction, with K_D_^app^ values of > 5 mM and > 1 mM, respectively (Figure 6f, e). Deletion of the COM^D^ region further reduced binding: T3-ΔCOM^D^_BpsA_/C3_BpsB_ yielded a K_D_^app^ of > 10 mM, whereas no measurable binding was observed for T6-ΔCOM^D^_BpsB_/C6_BpsC_ (Figure 6b, d).

**Figure 6.**
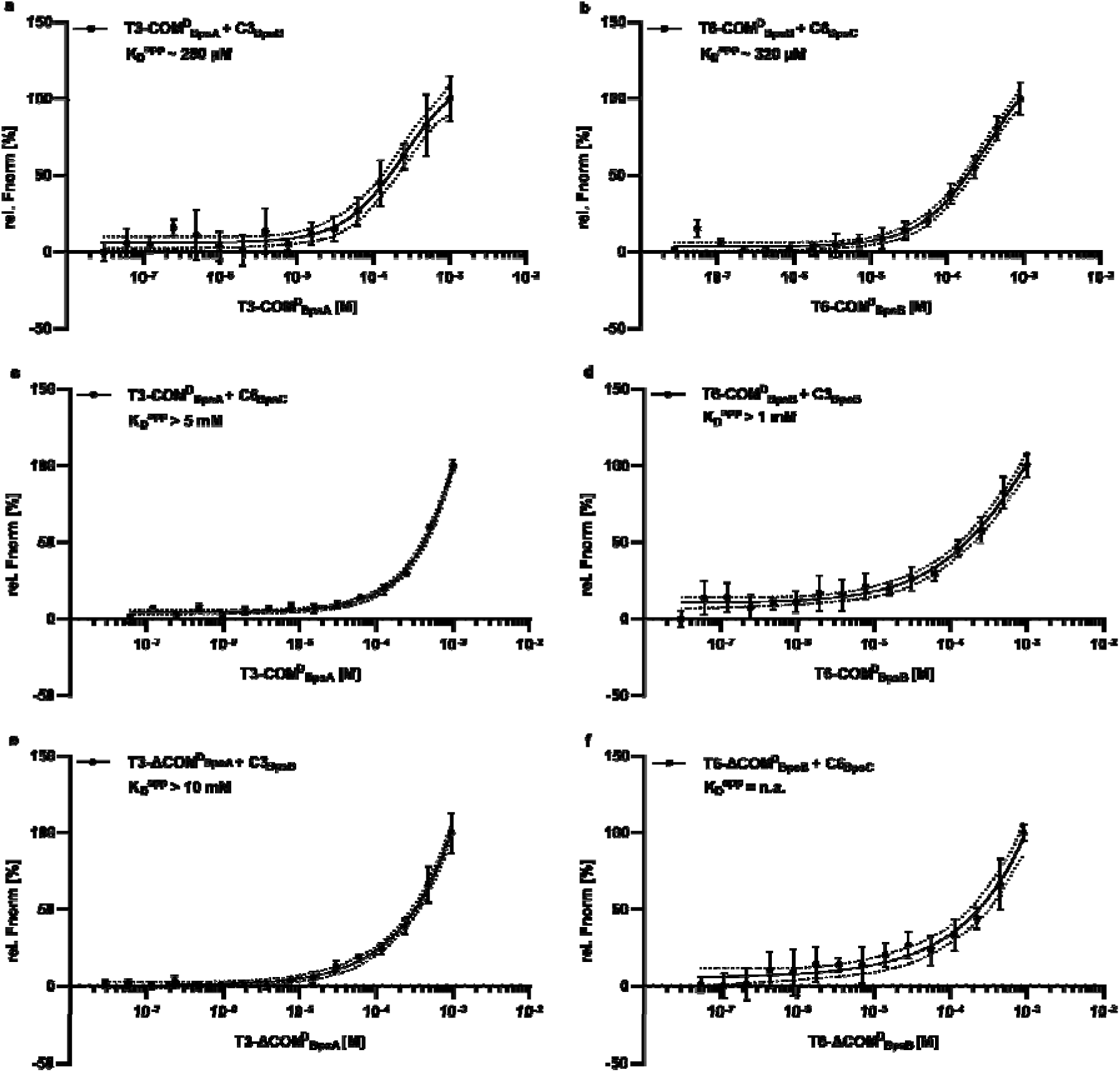
MST evaluation of *trans*-COM domain interactions between T3-COM^D^_BpsA_ and C3_BpsB_ (a) as well as T6-COM^D^_BpsB_ and C6_BpsC_ (b), possible crosstalks between non-adjacent *trans*-COM domain pairs within the NRPS of *A. balhimycina* (c,d) and constructs lacking the COM^D^ domains serving as controls (e,f). All data are plotted in relative, normalized fluorescence [%] against the ligand concentration [M] in a log_10_ scale,). Measurement was done in triplicate with the confidence level set to 95% (dotted line), Error bars indicate standard deviation (SD). MST measurements: MST power = medium, excitation power = 20-100% (auto-detect).

The reduced interaction observed for non-cognate combinations indicates that these donor-acceptor interactions are selective. Furthermore, the marked loss of binding upon deletion of the COM donor region demonstrates that this region is a major determinant of intermodular recognition. Together, these data indicate that COM^D^ regions contribute substantially to selective recognition and are important for intermodular binding.

### All internal C domains of BpsA and BpsB contain *cis*-COM domains

To assess whether the COM^A^ region structural motif identified in the N-terminal C domains of BpsB and BpsC is conserved across other domains, we predicted and compared all C domains and related homologous domains within BpsA, BpsB, and BpsC. Domain annotation using InterProScan, identified two C domains and one E domain in BpsA, three C domains and two E domains in BpsB, and one C domain together with a terminal X domain in BpsC (Figure 7). AF3 predictions yielded high-confidence structures for all ten domains, with pTM values ranging from 0.85 to 0.92 and consistently high pLDDT scores, supporting the reliability of the models.

**Figure 7.**
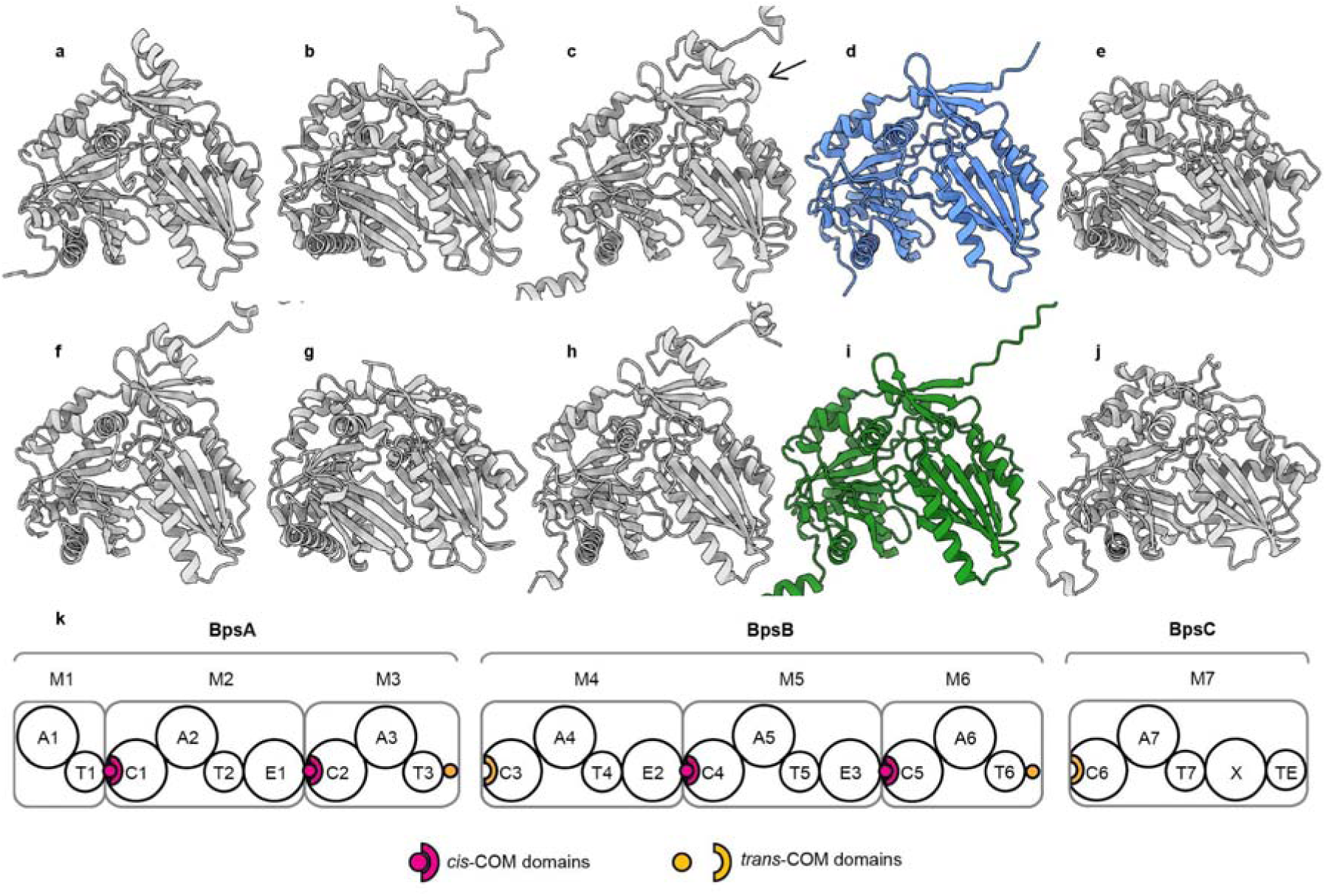
Predicted structures of all six C domains and the E- and X-domains from BpsA, BpsB, and BpsC and their location in the NRPSs. The two C domains of BpsA (**a**, **c**) and the second and third C domain of BpsB (**f**, **h**) show a N-terminal helix, which does not exist in the first C domain of BpsB (**d**) and the first C domain of BpsC (**i**). This helix is located in the same region of the C domains as the C-terminal helix of BpsA and BpsB when bound to the first C domains of BpsB and BpsC (Figure 3a, **b**). The arrow highlights the discussed region in the second C domain of BpsA, which differs from the other three internal C domains (see main text). A structural motif similar to the COM^A^ region in C domains can also be found in the E domain of BpsA (**b**) and the two E-domains of BpsB (**e**, **g**). In the X-domain of BpsC (**j**) this structural motif is barely visible. (All domains are depicted in the same orientation and colors are as before). (**k**) Schematic representation of the modular balhimycin NRPSs BpsA-C. Dots and semicircles in dark pink indicate the location of *cis*-COM domains and in yellow the location of *trans*-COM domains. A, adenylation domain; T, thiolation domain; C, condensation domain; E, epimerization domain; X, X domain; TE, thioesterase domain; M, module. Numbers indicate the domain order.

All six predicted C domains display the structural motif of a COM^A^ region, comprising a three stranded antiparallel β-sheet, an adjacent α-helix, and a loop (Figure 7a, c, d, f, h, i). In the N-terminal C domains of BpsB and BpsC (Figure 7d, i), this motif forms the interface that binds the C-terminal COM^D^ region of BpsA or BpsB, respectively. In contrast, in the other four internal C domains (Figure 7a, c, f, h), this motif interacts with a helix connected to the first strand of the β-sheet. This arrangement corresponds to a *cis*-COM domain.

Among the internal C domains, only the second C domain of BpsA exhibits a slightly longer connecting segment, consisting of two residues plus a three-residue α-helix (Figure 7c), while the remaining three C domains have a short three-residue connection (Figure 7a, f, h). This variation may reflect the evolutionary history of the system, potentially arising from gene fusion events that combined previously separate NRPS modules into BpsA^19^.

The three E domains share partial structural similarity with the COM^A^ structural motif, showing elements of the β-sheet and the adjacent α-helix and loop (Figure 7b, e, g). In contrast, the X domain lacks the β-sheet entirely (Figure 7j). Neither the E domains nor the X domain display a connected α-helix characteristic of *cis*-COM domains.

### *Cis*- and *trans*-COM domains are present in NRPSs from all five GPA classes

To determine whether cis- and trans-COM domains are conserved across NRPSs of the other GPA classes, we examined NRPS systems from GPA type II-V. Specifically, we analyzed the NRPSs from *Actinoplanes teichomyceticus* ATCC 31121, which synthesizes the type IV GPA teicoplanin^20,21^; *Amycolatopsis japonica* DSM 44213, which produces the type III GPA ristocetin^22^; *Amycolatopsis keratiniphila* subsp. *nogabecina,* the producer of the type II GPA actinoidin^23,24^; and *Nonomuraea* sp. ATCC55076, which produces the type V GPA kistamicin^25^. We first predicted the structures of all C domains in these NRPSs to assess the presence of *cis*-COM domains and then examined potential *trans*-COM domains in more detail.

In contrast to balhimycin, each of the above-mentioned GPAs is synthesized by four NRPSs, containing a total of six C domains. The first, fourth, and fifth C domains are internally located, whereas the second, third, and sixth C domains are N-terminal. The predicted structures of all C domains (Figure S3-6) showed high pTM values ranging from 0.84 to 0.91, indicating a correctly predicted overall fold. The pLDDT scores were mostly high (>70) to very high (>90) in the relevant regions of the C domains, indicating high confidence predictions. All predicted structures of the internal C domains displayed the structural motif characteristic of a *cis*-COM domain. In contrast, the predicted structures of the N-terminal C domains all lacked the helix that forms part of the *cis*-COM domain structural motif.

To further analyze the N-terminal C domains, we predicted their structures together with 22 to 25 C-terminal amino acids of their preceding NRPSs. For the third and sixth C domain of the kistamicin NRPS assembly line from *Nonomuraea* sp. ATCC55076, we instead used the amino acids spanning from the beginning of the C-terminal T domain to the C-terminus of the preceding NRPS. The predicted structures showed good pTM values between 0.79 and 0.87, again suggesting a correct overall fold of the predictions. The pLDDT scores were mostly high (>70) to very high (>90) in the relevant regions, thereby indicating high confidence predictions. For each predicted combination, at least one structure had an ipTM >0.8. Overall, the ipTM values ranged from 0.81 to 0,88, indicating high confidence in the predicted interfaces. All twelve predicted interfaces clearly exhibited the structural motif of a *trans*-COM domain (Figure S7, 8). Among these twelve *trans*-COM domains, the one involving the sixth C domain of the kistamicin NRPS assembly line from *Nonomuraea* sp. ATCC55076 differed most strongly from the others. In this case, the helix in the COM^D^ region is followed by a 25-amino-acids-long disordered region.

Collectively, these structural predictions demonstrate that *cis*- and *trans*-COM domains occur in NRPSs from all five GPA classes.

## Discussion

In GPA-producing NRPS assembly lines, successive modules must coordinate substrate transfer with high precision, whether they are located within the same protein or distributed across separate proteins. However, the molecular basis of this intermodular coordination remains incompletely understood. Here, we combine genetic perturbation, structure prediction and quantitative interaction analyses to characterize how *trans*-COM domains mediate intermodular interactions in the balhimycin NRPS assembly line. We further identify *cis*- and *trans*-COM domains at each module-module junction across GPA-producing NRPS assembly lines representing all five GPA classes.

Our *in vivo* data show that the COM^D^ regions in BpsA and BpsB are required for efficient balhimycin biosynthesis, although they are not strictly essential. The strong decrease in balhimycin production observed upon deletion of the COM^D^ regions suggests that COM domain-mediated interaction enhances the efficiency of intermodular transfer rather than serving as an absolute prerequisite for catalysis. The remaining residual production might be explained by the interaction of the T domains and the respective C domains.

The AF3 structure predictions, together with the MST measurements, revealed that the two identified *trans*-COM domain pairs in the balhimycin assembly line interact specifically but with relatively low affinity. Notably, under our assay conditions, the measured affinities were weaker than those reported for several previously characterized COM domains^15,17^ or docking domains^26–30^. One possible explanation is the T-COM^D^/C domain architecture of the *trans*-COM domains in the balhimycin NRPS assembly line. In this arrangement, the COM^D^ region is not connected to an upstream E domain as in the E-COM^D^/C topology, for which different affinities were reported^15,17^. Here, the *trans*-COM domains are proposed to stabilize the E-C domain orientation through a scaffolding role. This suggests that *trans*-COM domains in a T-COM^D^/C architecture either lack this scaffolding role or perform it differently, potentially reducing the requirement for higher-affinity interactions.

Our comparative analysis across GPA biosynthetic systems identified *cis*- or *trans*-COM domains upstream of every C domain in the five GPA NRPS assembly lines investigated. Among these, we identified *cis*-COM domains in a previously undescribed T-COM–C domain topology (Figure 6k). As in the T-COM/C *trans*-COM arrangement, no E domain is located upstream of the COM domain in this topology. These *cis*-COM domains are therefore unlikely to mediate the E-C scaffolding function proposed for E-COM/C architectures.

Instead, their structural conservation across phylogenetically and biosynthetically distinct GPA systems suggests broader roles in intermodular communication, acceptor T domain recruitment, or substrate handover.

The structural similarities between *cis*- and *trans*-COM domains, together with their appearance at each module-module junction across the analyzed GPA-producing NRPS assembly lines, support the idea that these domains represent mechanistically related solutions for organizing NRPS assembly lines.

Comparison of our high-confidence AF3-predicted structures of *cis*- and *trans*-COM domains with recently reported cryo-EM^16^ and crystal^17^ structures indicates that AF3 is a suitable tool for COM domain analysis. Although predictions of other NRPS interdomain interactions, such as those between T and C domains, remain more challenging for AF3, the predicted COM domain structures and associated confidence scores are highly consistent with recently reported experimentally determined structures.

Together, our results suggest that COM domains act as versatile and conserved intermodular coordination elements in GPA NRPS assembly lines. Depending on their domain topology and molecular context, they may contribute to docking, partner selection, acceptor T domain recruitment, substrate-transfer coordination and structural scaffolding. Nevertheless, the limited number of GPA NRPS assembly lines investigated here highlights the need for further systematic structural and functional characterization. These findings broaden our understanding of COM domains in GPA biosynthesis and provide a conceptual basis for the targeted engineering of NRPS assembly lines, with the long-term goal of generating new GPAs.

## Resource availability

### Lead contact

Further information and requests for resources and reagents should be directed to and will be fulfilled by the lead contact, Evi Stegmann

### Materials availability

This study did not generate new unique reagents.

### Data and code availability

The data supporting the findings of this study are available within the article and its supplementary information.

- The predicted AF3 models have been deposited in the Zenodo repository under accession number #####.
- This study did not generate any unique code.
- Any additional information required to reanalyze the data reported in this paper is available from the lead contact upon request.

## Acknowledgments

The work related to this study was conducted in the laboratories of E.S., W.W., and T.S. was supported by the Deutsche Forschungsgemeinschaft (DFG, German Research Foundation) through the Collaborative Research Centre/Transregio TRR 261 "Cellular Mechanisms of Antibiotic Action and Production" (Projects B01 and B08). We thank Athina Gavriilidou and Nadine Ziemert for providing initial help in the identification of the COM domain sequences and Libera Lo Presti for valuable comments on the manuscript.

## Author contributions

ES, TS, and WW conceived the study. T.M.H. performed the *in silico* modeling and analyzed the resulting data. D.B. conducted the *in vivo* experiments and analyzed the resulting data. N.P. performed the in vitro interaction experiments and analyzed the resulting data. A.K. carried out the HPLC–MS analyses. D.B., T.M.H., and E.S. drafted and wrote the original manuscript. All authors reviewed, revised, and approved the final version of the manuscript.

## Supplemental information

Supplementary data: Figures S1-8 & Tables S1-4,

## Methods

### Bacterial strains, cultivation conditions and molecular cloning

Cultivation and DNA manipulation in *Escherichia coli* NovaBlue were performed as previously described for *E. coli* ^31^. For plasmid selection, the medium was supplemented with apramycin (100 µg/ml), ampicillin (150 µg/ml), or kanamycin 50 µg/ml. *Amycolatopsis balhimycina* and all derived mutants were cultivated at 29°C, in baffled flasks with a steel coil and shaken at 120 rpm in R5 medium^32^ or on solid agar (1.5% (w/v)) medium. Apramycin 50 μg/ml or erythromycin 50 µg/ml was used for plasmid selection.

Oligonucleotides used for mutant constructions are listed in Table S2. All used strains and plasmids are listed in Table S3 and S4, respectively.

For all polymerase chain reactions (PCRs), the “Q5 High-Fidelity DNA Polymerase (NEB©)” was used to amplify the genes with the overhangs for the respective target vector. All enzymes were from Thermofischer. For deletions in *A. balhimycina* the pSP1 vector was linearized using the restriction endonuclease *Kpn*I. DNA fragments were assembled by Gibson assembly^33^ using the NEBuilder HiFi DNA assembly Mix (NEB©). DNA gel extraction and plasmid isolation were performed using the purification kits from Macherey-Nagel (#REF 740609.50) and VWR (#13-6945-00).

### Genetic manipulation of *A. balhimycina*

Genetic manipulation of *A. balhimycina* in this study was achieved using a well-established direct transformation protocol.^3^ This requires the isolation of demethylated DNA from the *E. coli* strains JM110 and ET12567. For the analysis of the COM domains, *in-frame* deletion mutants of *A. balhimycina* were generated, in which the C-terminal donor COM domain encoding region was deleted using a homologous recombination approach. Deletion was performed using pSP1^3^ derivatives containing flanking homologous regions. The flanking regions of approximately 1,5 kB up- and downstream of the COM domains were cloned using primers P1-P4 and P10-P13 (Table S2) and the corresponding plasmids were integrated by direct transformation. Erythromycin-resistant clones harboring pSP1ΔDB1, or pSP1ΔDB2 on the chromosome, were confirmed by PCR using primers P5-P6 (Table S2) and subsequently used in the stress protocol to induce a crossover event according to Puk et al.^34^. The generated protoplasts were streaked on R5-agar plates as well as R5-agar plates supplemented with 50 µg/ml erythromycin. Erythromycin-sensitive clones were screened for in-frame deletion by PCR using primers P7-P8 and P19-P20 (Table S2).

### Balhimycin production and HPLC-MS analysis

For the production assay, the generated mutants and the wildtype strain were inoculated in 20 ml R5-Medium and incubated for 48h as preculture as described above. After growth, 5 ml were taken to inoculate 100 ml of R5-Medium as the main culture. After 120h of cultivation, 10 ml of cultures were harvested and separated by centrifugation into supernatant and mycelium. The supernatant was directly used for bioactivity assays as well as for HPLC-MS measurements. The mycelium was lyophilized and weighed for dry cell weight determination and subsequent balhimycin quantification.

Balhimycin detection was performed by analyzing 2.5 μl of the different supernatants by means of high-performance liquid chromatography-electrospray ionization-mass spectrometry (HPLC-ESI-MS) using a Nucleosil 100-C18 column (3 μm, 100 by 2 mm) (precolumn, 10 by 2 mm) (Dr. Maisch GmbH, Ammerbuch-Entringen, Germany) coupled to an ESI mass spectrometer. LC-MS measurements were obtained from a LC/MSD Ultra Trap system XCT 6330 (Agilent Technologies, Waldbronn, Germany). Analysis was carried out at a flow rate of 400 µl/min with gradient elution. Solvent A was 0.1% formic acid, and solvent B 0.06% formic acid in acetonitrile. Gradient elution was performed as follows: t0=0% B, t15=t17=100% B, post time 5 min. 0% B. The flow rate was 400 µl/min, and the temperature was 40°C. For MS analysis, an electrospray ionization (alternating positive and negative ionization) in Ultra Scan mode with a capillary voltage of 3.5 kV and a drying gas temperature of 350°C was used. Detection of m/z values was conducted with Agilent DataAnalysis for 6300 series Ion Trap LC/MS 6.1 version 3.4 software (Bruker-Daltonik GmbH).

To quantify the amounts of balhimycin produced by the mutants the peak intensities of each data point in the MS chromatogram corresponding to the masses of balhimycin were summed to a total intensity. This was used to determine the concentration of balhimycin by comparing it to a calibration curve generated by measuring pure balhimycin. To compare the produced amounts, all production levels of balhimycin were normalized to the cell dry weight.

### MicroScale Thermophoresis (MST)

To perform MST measurements, *trans*-COM domain constructs with the respective predicted sequence and one adjacent domain were included in the construct design. For BpsA, the C-terminal T-domain was attached to the subsequent COM^D^ region, resulting in a T3-COM^D^_BpsA_ construct. The same was done for T6-COM^D^_BpsB_, consisting of the C-terminal T-domain of BpsB and the respective COM^D^ region. C3_BpsB_ and C6_BpsC_ comprised the N-terminus through the end of the first C domain of BpsB and BpsC, respectively, including the COM^A^ regions. For control experiments, C-terminal *trans*-COM domains were excluded, giving a construct containing the adjacent T domain. For all MST measurements, the RED – MALEIMIDE 2nd Generation Kit (NanoTemper) was used. The coupling of the fluorophore to the target protein was done as described in the manual appended to the Kit. All measurements were performed at room temperature using the Monolith NT.115 NanoTemper MST device with the instrument settings: MST power = medium and excitation power = 100% (auto-detect).

MST buffer (25 mM HEPES/NaOH, 50 mM NaCl, pH 7.5) was used to dilute the target protein prior to the dilution series. Maleimide labelling buffer (NanoTemper) was used to perform the dilution series prior to the actual measurement. The unlabelled ligand, in this case the binding protein, was concentrated to at least 2 mM for all experiments. The concentration of the labelled target protein was adjusted according to the labelling efficiency (200 – 400 nM). All samples were centrifuged at max. speed for 10 min prior use to sediment possible aggregates. For the measurement, Monolith NT.115 Premium Capillaries were used (NanoTemper) to avoid adsorption of the molecules to the inner surface of the capillaries. All measurements were done in triplicate. Data points were initially normalized and then fitted using a sigmoidal, 4PL fit (GraphPad).

### Prediction of protein-protein interfaces

Protein-protein interactions were predicted with AlphaFold3 (AF3) as implemented in AlphaFold Server^35^. The accuracy of the predictions was assessed using the confidence metrics provided by AF3, namely the predicted Template Modeling score (pTM), the interface predicted Template Modeling score (ipTM), the predicted Local Distance Difference Test (pLDDT), and the Predicted Aligned Error (PAE).

A pTM score greater than 0.5 suggests that the overall fold of the predicted complex may resemble the native structure. ipTM scores above 0.8 indicate confident, high-quality predictions, whereas scores below 0.6 suggest that the predicted interface is likely incorrect. Scores between 0.6 and 0.8 represent an intermediate-confidence range in which predictions may be either correct or incorrect.

The pLDDT score provides a per-atom confidence estimate ranging from 0 to 100. Scores above 90 indicate very high confidence, scores between 70 and 90 indicate high confidence, scores between 50 and 70 indicate low confidence, and scores below 50 indicate very low confidence.

PAE estimates the expected error in the relative positions and orientations of pairs of tokens within the predicted structure. Higher PAE values correspond to greater predicted errors and, consequently, lower confidence in the relative placement of the respective tokens.

All molecular visualizations were created with UCSF ChimeraX (v 1.9)^36^.

### Analysis of the predicted protein-protein interfaces

The primary forces involved in protein-protein interactions are hydrophobic interactions, electrostatic interactions, hydrogen bonds, and Van der Waals forces. To check the plausibility of the AF3 predicted protein-protein interactions the molecular lipophilicity potential, the electrostatic potential, the number of salt bridges and hydrogen bonds, and the area of the interface were calculated and analyzed. The molecular lipophilicity potential was investigated with UCSF ChimeraX using the command mlp^36–39^.

APBS (v 3.4.1)^40^ was used to calculate the electrostatic potentials of the interacting proteins. As part of these calculations PDB2PQR (v 3.6.1) and PROPKA (v 3.5.1)^41,42^ were used. The protonation states of the proteins were determined at a pH of 7 and 7.5 with PROPKA. For all proteins the protonation states at pH 7 and 7.5 were identical. The electrostatic potentials were calculated at an ionic strength of 150 mM (NaCl) and a temperature of 298.15 K. As values for the dielectric constants 2 and 78.54 were used for the protein and the solvent respectively.

To identify the amino acids involved in the predicted protein-protein interfaces, to evaluate the number of salt bridges and hydrogen bonds and to determine the area of the protein interface PISA (v 1.52) was used as implemented at the European Bioinformatics Institute^43^.

For the annotation of domains InterProScan was used as implemented at the European Bioinformatics Institute^44,45^. The annotation of the NRPSs was done with antiSMASH^46^.

