## Supplementary data for "Conserved *cis* and *trans* communication domains mediate module interaction in glycopeptide antibiotic NRPS assembly lines"

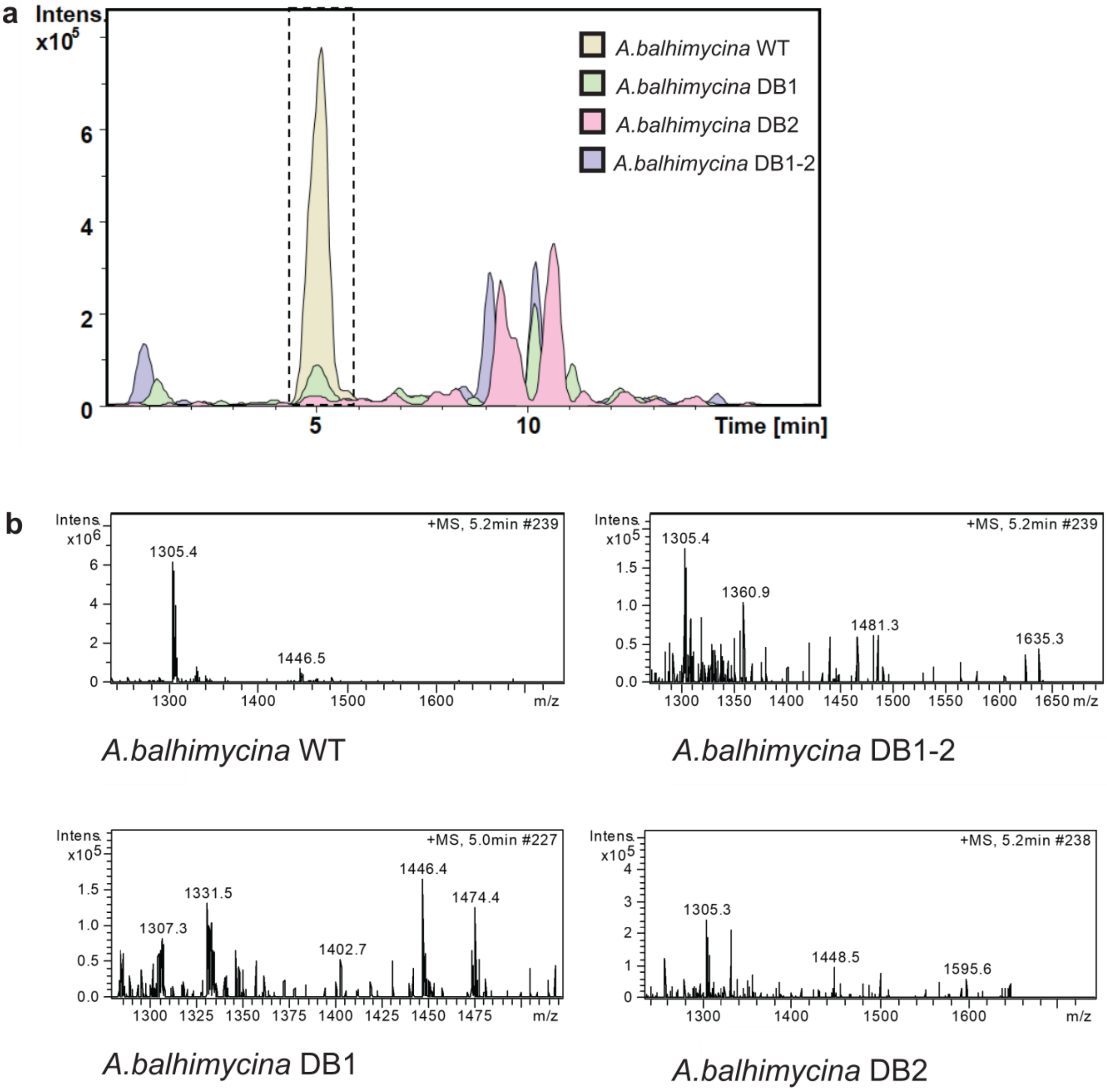


**Figure S1|HPLC-MS analysis of the *A. balhimycina* DD mutants.** (**a**) Chromatograms showing the peak for balhimycin detection (RT) and (**b**) the corresponding MS spectra of each chromatogram displaying the masses of balhimycin.

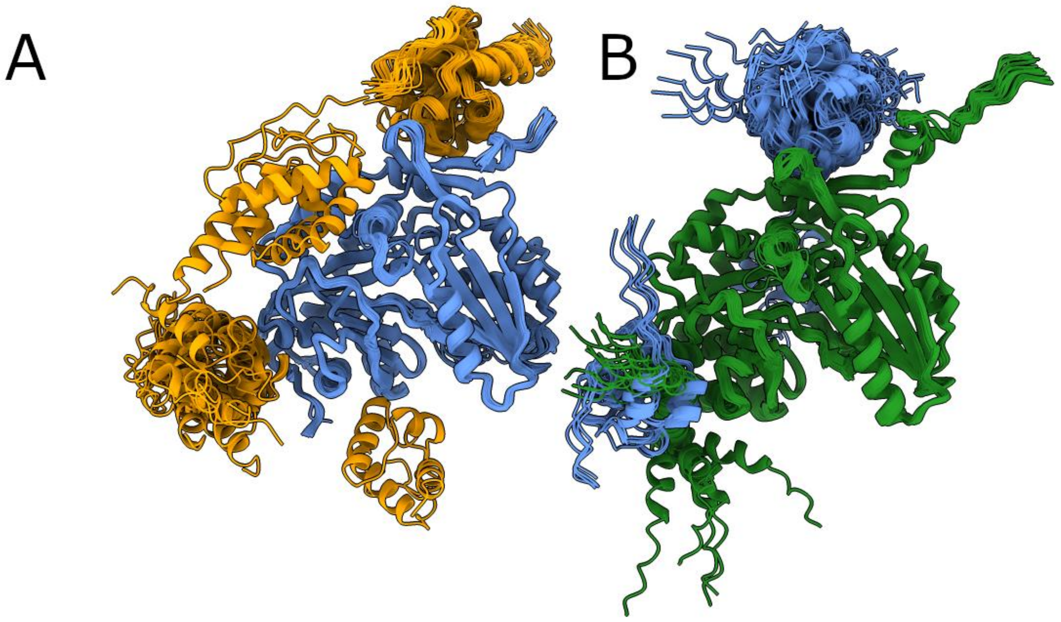


**Figure S2|Interactions of the C-terminal T domain of BpsA (orange) and BpsB (blue) with the N-terminal C domain of BpsB and BpsC (green) as predicted by AF3.** The 25 superimposed models of predicted interactions between the C-terminal T domain of BpsA and the N-terminal C domain of BpsB (**A**) show roughly four different relative orientations of the two domains. Something similar can be seen in the depiction of the 25 superimposed models of predicted interactions between the C-terminal T domain of BpsB and the N-terminal C domain of BpsC (**B**). Here three different relative orientations are visible. (All C domains are depicted in the same orientation and colors as before).

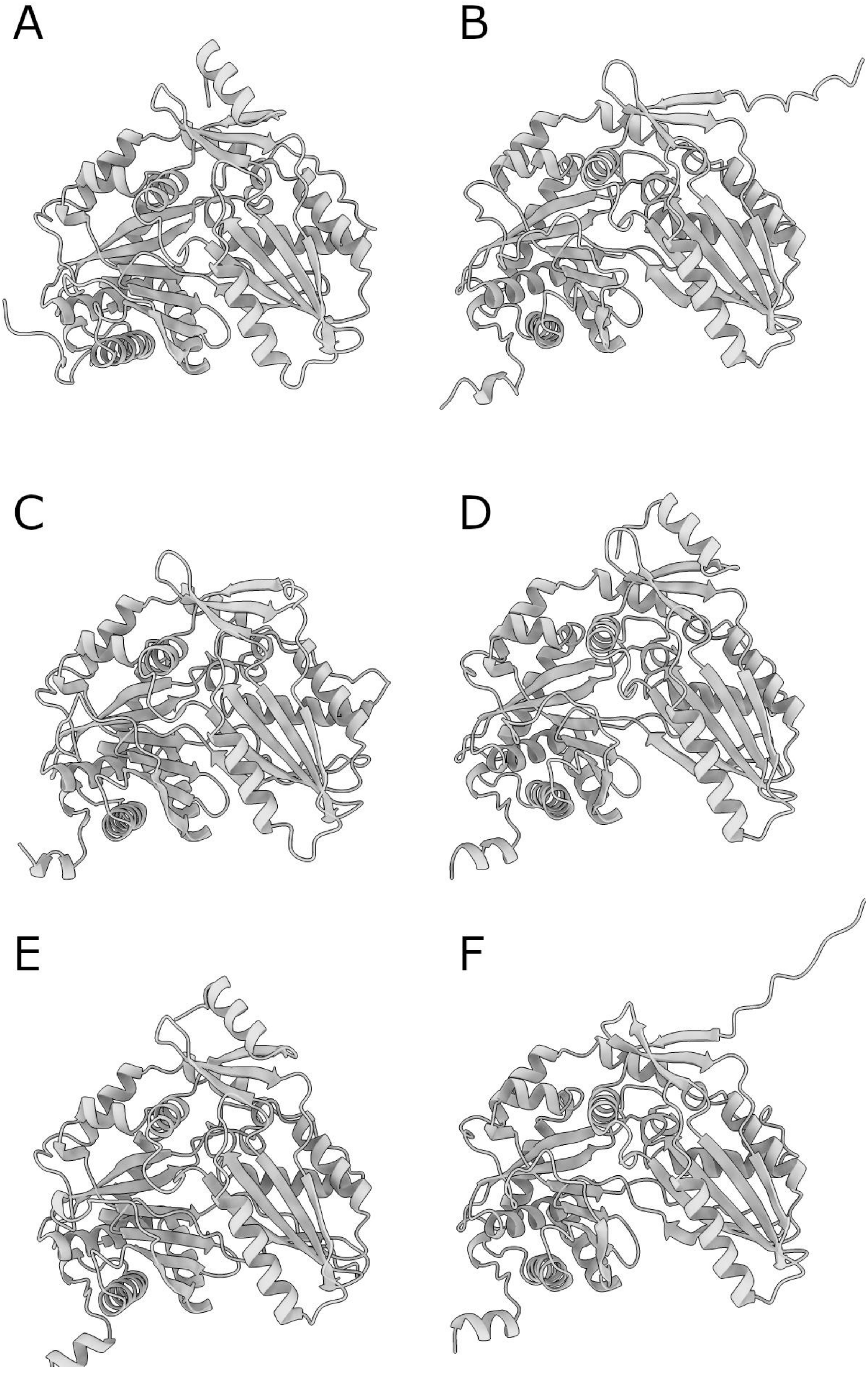
A
B
C
D
E
F

**Figure S3|Predicted structures of the six C domains from the four NRPSs of the teicoplanin biosynthetic gene cluster in *Actinoplanes teichomyceticus* ATCC 31121.** Teicoplanin is a type IV GPA. Out of the three internal C domains, one is located within the first (**A**) and two within the third NRPS (**D**, **E**). They show the structural motif of a *cis*-COM domain. The N-terminal C domains (**B**, **C**, and **F**) reside at the start of the second, third, and fourth NRPS, respectively, and display the $COM^{A}$ region characteristic of a *trans*-COM domain.

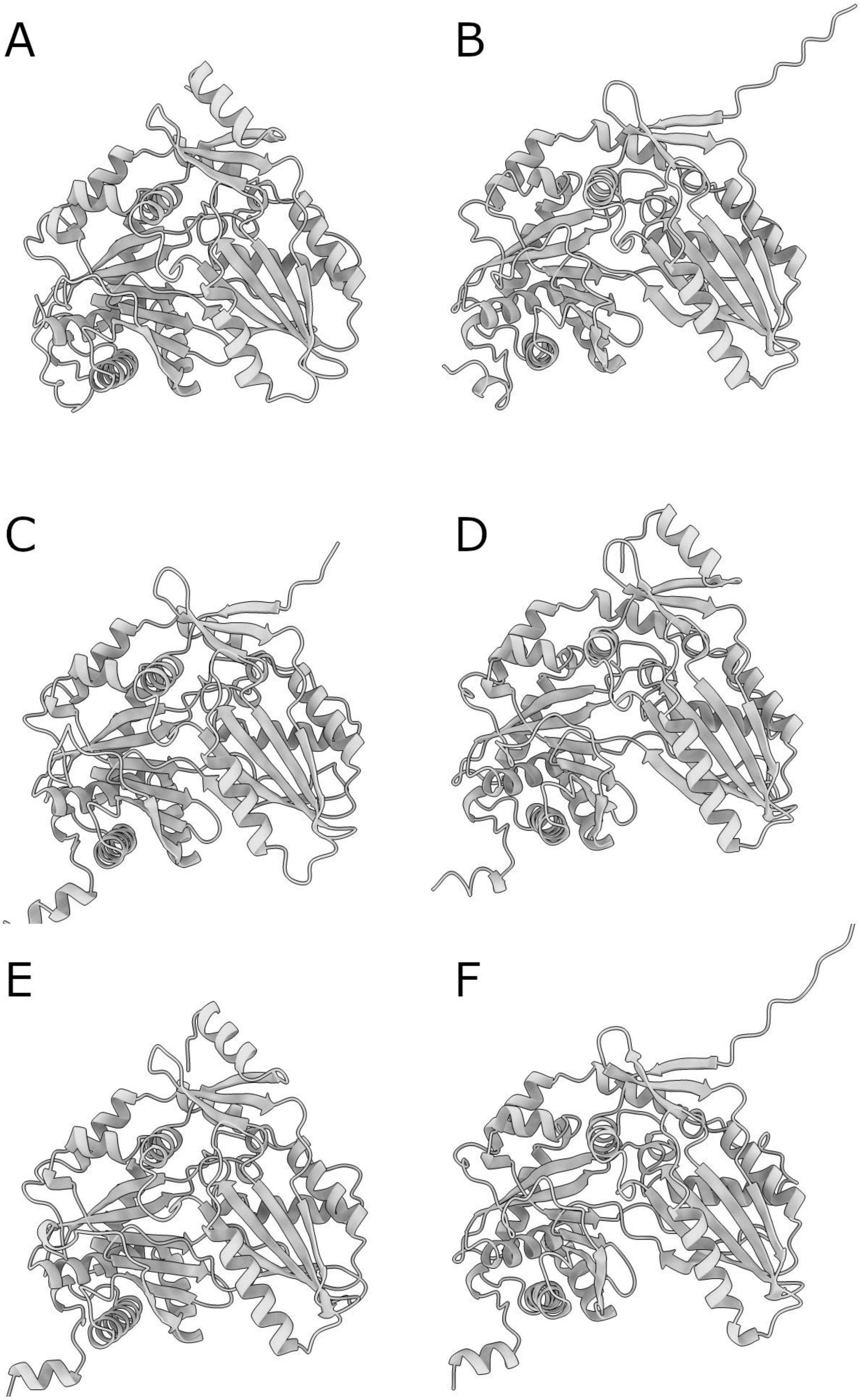
A
B
C
D
E
F

**Figure S4|Predicted structures of the six C domains from the four NRPSs of the ristocetin biosynthetic gene cluster of *Amycolatopsis japonica* DSM 44213.** Ristocetin, a type III GPA, is synthesized by four NRPSs harboring six C domains. These include an internal C domain from the first NRPS (**A**); an N-terminal C domain from the second NRPS (**B**); an N-terminal (**C**) and two internal C domains (**D**, **E**) from the third NRPS; and an N-terminal C domain from the fourth NRPS (**F**). The internal C domains (**A**, **D**, and **E**) exhibit the *cis*-COM structural motif, while the N-terminal C domains (**B**, **C**, and **F**) display the $COM^A$ region of the *trans*-COM domain.

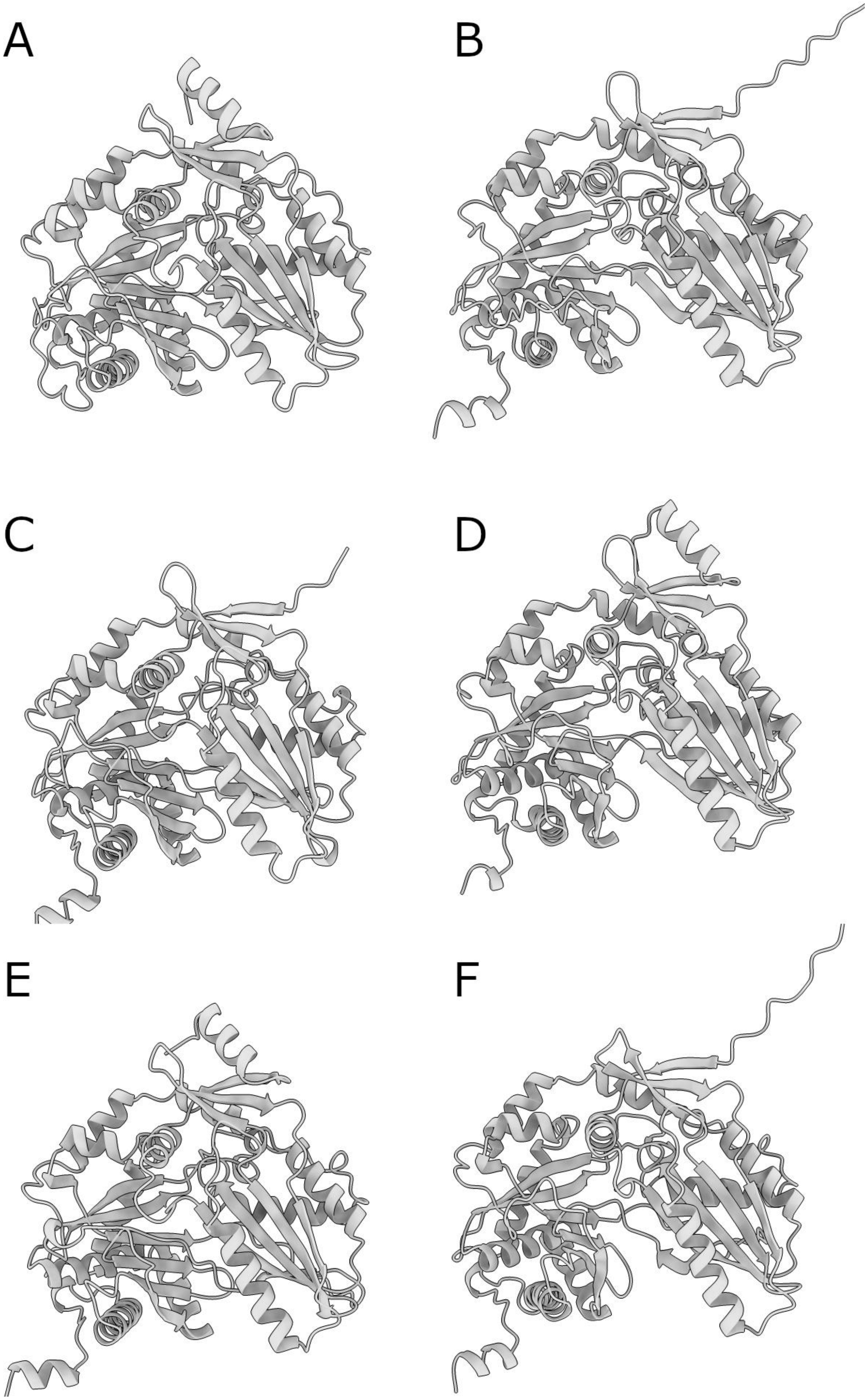
A
B
C
D
E
F

**Figure S5|Predicted structures of the six C domains from the four NRPSs of the actinoidin biosynthetic gene cluster from *Amycolatopsis keratiniphila* subsp. *nogabecina*.** Actinoidin, a type II GPA, is synthesized by four NRPSs containing six C domains. These include internal C domains from the first (**A**) and third (**D**, **E**) NRPS, as well as N-terminal C domains from the second (**B**), third (**C**), and fourth (**F**) NRPS. The internal C domains (**A**, **D**, and **E**) show a *cis*-COM structural motif, while the N-terminal C domains (**B**, **C**, and **F**) display the structural motif of a $COM^{A}$ region of a *trans*-COM domain.

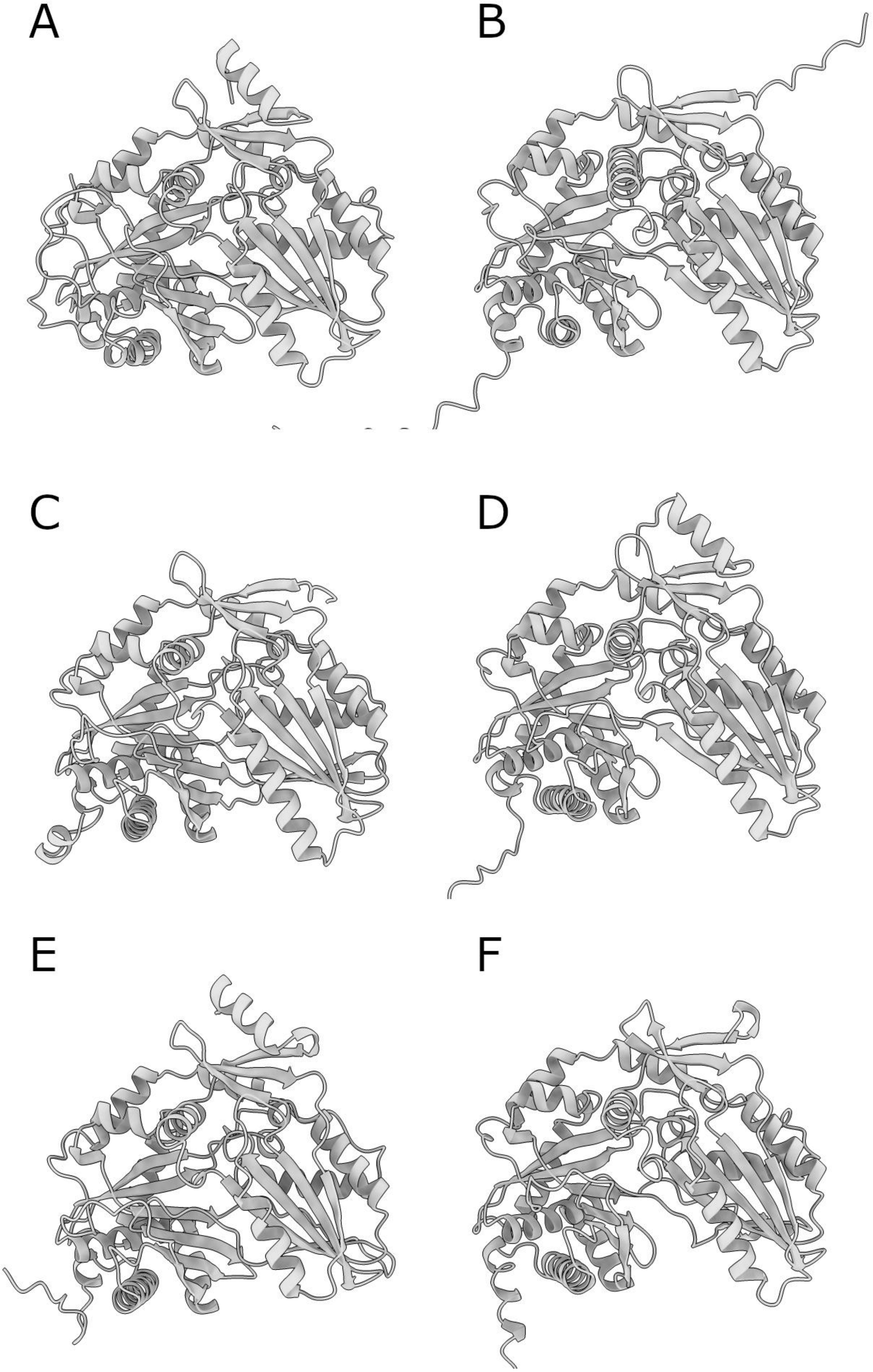
A
B
C
D
E
F

**Figure S6|Predicted structures of the six C domains from the four NRPSs of the kistamycin biosynthetic gene cluster from *Nonomuraea* sp. ATCC55076.** Kistamycin is a type V GPA synthesized by four NRPSs containing six C domains. Among them one internal C domain from the first (**A**) and two from the third (**D**, **E**) NRPS, and one N-terminal C domain from the second (**B**), third (**C**), and fourth (**F**) NRPS respectively. The internal C domains (**A**, **D**, and **E**) feature the structural motif of a *cis*-COM domain, while the N-terminal domains (**B**, **C**, and **F**) exhibit the $COM^A$ region of a *trans*-COM domain.

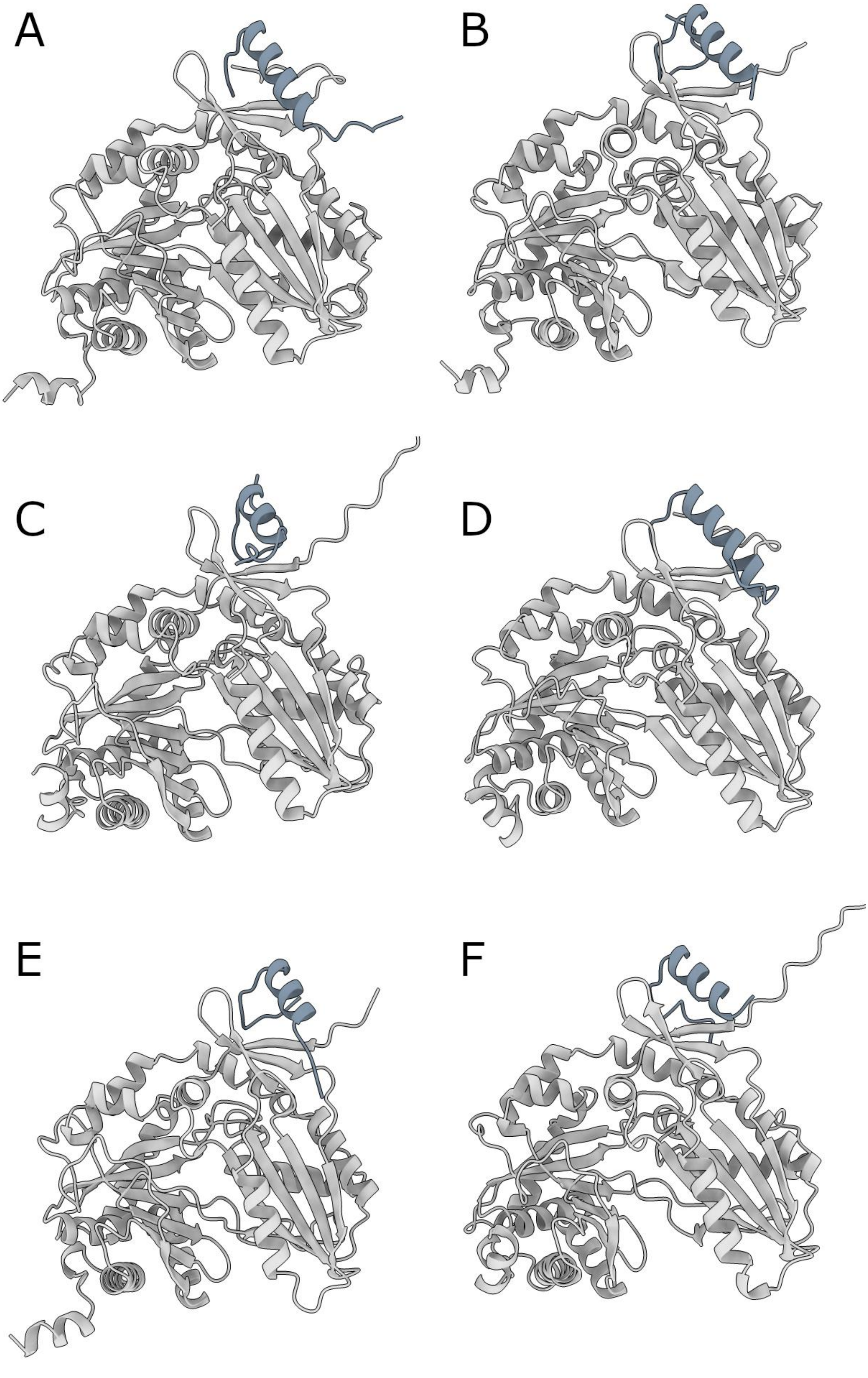
A
B
C
D
E
F

**Figure S7|Predicted structures of the *trans*-COM domains from *Actinoplanes teichomyceticus* ATCC 31121 and *Amycolatopsis japonica* DSM 44213.** The three *trans*-COM domains of *A. teichomyceticus* ATCC 31121 involve the second (**A**), third (**B**), and sixth (**C**) C domain. In *A. japonica* DSM 44213 also the second (**D**), third (**E**), and sixth (**F**) C domains comprise trans-COM regions. Across all models, the *trans*-COM$^{D}$ regions are rendered in grey, while the C domains encompassing the *trans*-COM$^{A}$ regions are depicted in silver.

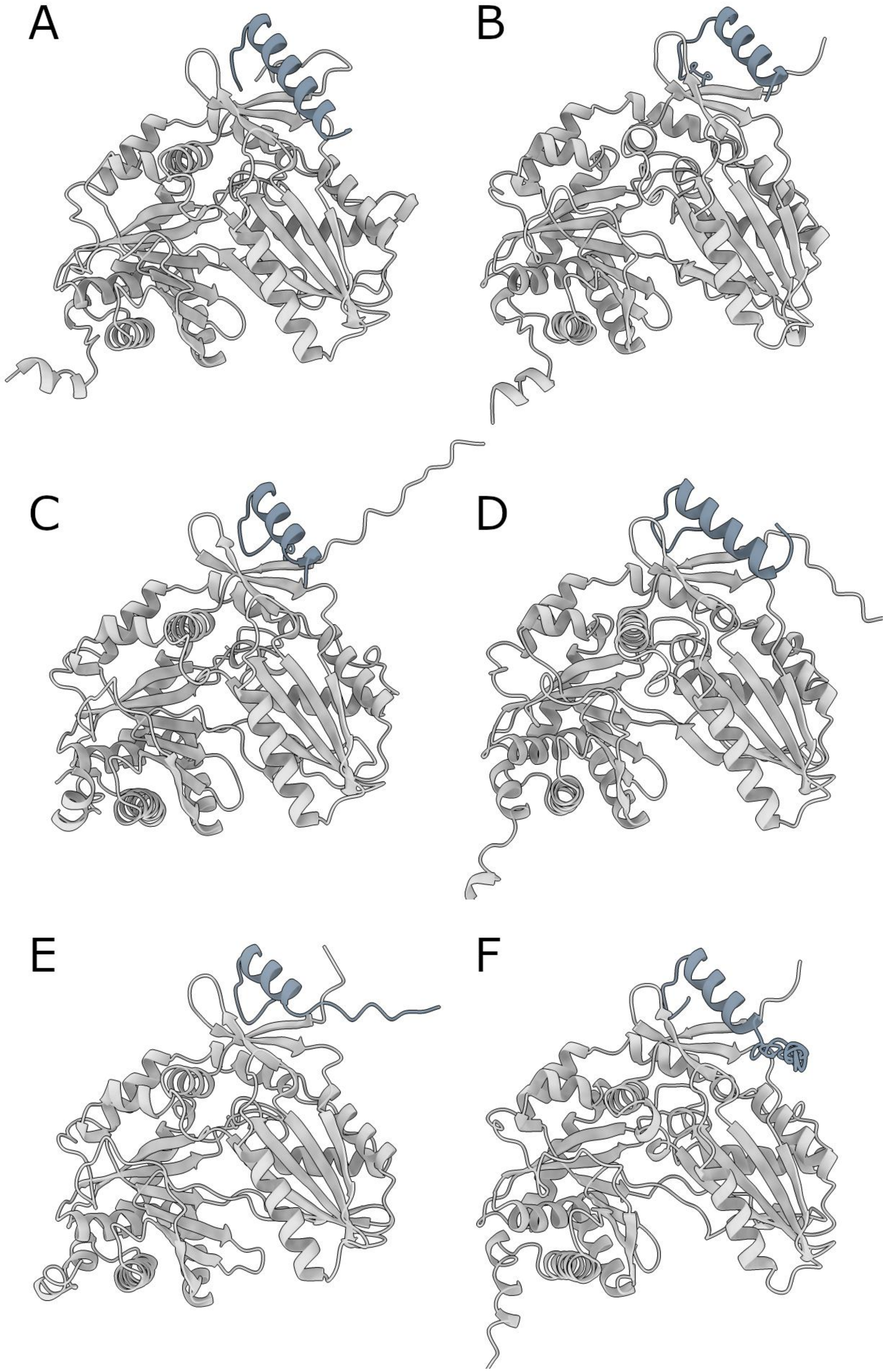
A
B
C
D
E
F

**Figure S8|Predicted structures of the *trans*-COM domains from *Amycolatopsis keratiniphila* subsp. *nogabecina* and *Nonomuraea* sp. ATCC55076. T**he three *trans*-COM domains of *A. keratiniphila* subsp. *nogabecina* (**A**, **B**, and **C**) and the three *trans*-COM domains of *Nonomuraea* sp. ATCC55076 (**D**, **E**, and **F**) are shown. The *trans*-$COM^D$ regions are depicted in grey, whereas the C domains, including the embedded *trans*-$COM^A$ regions, are shown in silver.

**Table S1|Description of the two interfaces between the α-helices and the C domains.** Results of a PISA interface analysis of 5 models from 5 different AF3 runs for each of the two interfaces. It lists the numbers of amino acids and atoms involved in the interface, the number of hydrogen bonds and salt bridges in the interface, and the interface area of the α-helix and the linker.

| nr | model description | #AAs / #atoms in interface ($COM^A$ \| $COM^D$) | #HB in interface | #SB in interface | Interface area (SAA) of $COM^D$ Å$^2$ (%) |
|---|---|---|---|---|---|
| **1** | $COM^D_{BpsA}$ and $COM^A_{BpsB}$ run 1 model 0 | 26/89 \| 16/72 | 9 | 9 | 881.9 (37.2) |
| **2** | $COM^D_{BpsA}$ and $COM^A_{BpsB}$ run 2 model 0 | 24/83 \| 15/67 | 11 | 10 | 844.1 (36.1) |
| **3** | $COM^D_{BpsA}$ and $COM^A_{BpsB}$ run 3 model 0 | 24/87 \| 16/72 | 13 | 11 | 868.6 (37.8) |
| **4** | $COM^D_{BpsA}$ and $COM^A_{BpsB}$ run 4 model 0 | 24/86 \| 16/70 | 12 | 10 | 881.2 (37.5) |
| **5** | $COM^D_{BpsA}$ and $COM^A_{BpsB}$ run 5 model 0 | 24/85 \| 15/68 | 12 | 11 | 854.3 (36.6) |
| | **average:** | **24.4/86 \| 15.6/69.8** | **11,4** | **10,2** | **866.02 (37.04)** |
| **6** | $COM^D_{BpsB}$ and $COM^A_{BpsC}$ run 1 model 0 | 24/74 \| 13/59 | 8 | 4 | 735.8 (34.9) |
| **7** | $COM^D_{BpsB}$ and $COM^A_{BpsC}$ run 2 model 0 | 24/75 \| 13/56 | 7 | 4 | 768.9 (36.4) |
| **8** | $COM^D_{BpsB}$ and $COM^A_{BpsC}$ run 3 model 0 | 24/74 \| 13/57 | 8 | 6 | 758.8 (35.8) |
| **9** | $COM^D_{BpsB}$ and $COM^A_{BpsC}$ run 4 model 0 | 24/72 \| 14/58 | 9 | 4 | 750.7 (35.5) |
| **10** | $COM^D_{BpsB}$ and $COM^A_{BpsC}$ run 5 model 0 | 24/72 \| 12/58 | 9 | 5 | 764.5 (36.2) |
| | **average:** | **24/73,4 \| 13/57.6** | **8,2** | **4,6** | **755.74 (35.76)** |

**Table S2|Oligonucleotides used in this study.**

| ID | Primer name | Sequence |
|---|---|---|
| P1 | cDDbpsB_UP_f | ggagagacgaattcgagctcggtacatggccgtgtcgttcgc |
| P2 | cDDbpsB_UP_r | cttatggcggtcattcctggacgatcaccgc |
| P3 | cDDbpsB_DO_f | atcgtccaggaatgaccgccataaggagcaga |
| P4 | cDDbpsB_DO_r | actcactatagggaaagcttgcatgggtcaggttcgcgccg |
| P5 | pSP1_lacZ_f | atgtgctgcaaggcgattaagt |
| P6 | pSP1_lacZ_r | gaaacagctatgaccatgattacgcc |
| P7 | KO_cDDbpsB_f | atgcagctgtcggcccg |
| P8 | KO_cDDbpsB_r | gcattccctcctgcagcg |
| P9 | KO_cDDbpsB_r_2 | agacgaacgagatccgctcggc |
| P10 | cDDbpsA_up_f | ggagagacgaattcgagctcggtactccgggctatccggcc |
| P11 | cDDbpsA_up_r | ccagccacgttcaccgttccacgatcgccg |
| P12 | cDDbpsA_down_f | atcgtggaacggtgaacgtggctggacatgttgt |
| P13 | cDDbpsA_down_r | actcactatagggaaagcttgcatgggcaccccattccccg |
| P14 | pSP1ΔcDDbpsA_seq1 | atcgccttcgatctgacggtcacc |
| P15 | pSP1ΔcDDbpsA_seq2 | agctggtgttcgccggccg |
| P16 | pSP1ΔcDDbpsA_seq3 | agcagcgcacgcccgcg |
| P17 | pSP1ΔcDDbpsA_seq4 | accgcctggtggtgacctgcc |
| P18 | pSP1ΔcDDbpsA_seq5 | acacgctgctcgtcttcgagaag |
| P19 | KO_cDDbpsA_f | agcggctgctgtgtgc |
| P20 | KO_cDDbpsA_r | gccagatttcctcgatccgcga |

**Table S3|Plasmids used in this study.**

| Plasmid | Relevant characteristics | Reference |
|---|---|---|
| pSP1 | pT7/T3-α19 derived gene disruption vector, with *ermE* gene in *Sap*I site EryR/AmpR | Pelzer S. et al.[1] |
| pSP1ΔDB1 | pSP1 derivative containing ~1,5kB homologous flanking regions for $COM^{D}_{BpsA}$ deletion | This study |
| pSP1ΔDB2 | pSP1 derivative containing ~1,5kB homologous flanking regions for $COM^{D}_{BpsB}$ deletion | This study |
| pET-24a(+) | T7 promoter, MCS(*Bam*HI-*Xho*I), His-Tag CDS, T7 terminator, lacI CDS, pBR322 ori, KanR, f1 ori | Novagen® |
| pET-24a(+)-T3-$COM^{D}_{BpsA}$ | pET-24a(+) backbone with T3-$COM^{D}_{BpsA}$ inserted in MCS with restriction sites *Nde*I/ *Bam*HI | This study |
| pET-24a(+)-T6-$COM^{D}_{BpsB}$ | pET-24a(+) backbone with T6-$COM^{D}_{BpsB}$ inserted in MCS with restriction sites *Nde*I/ *Bam*HI | This study |
| pET-24a(+)-T3-Δ$COM^{D}_{BpsA}$ | pET-24a(+) backbone with T3-Δ$COM^{D}_{BpsA}$ inserted in MCS with restriction sites *Nde*I/ *Bam*H I | This study |
| pET-24a(+)-T6-Δ$COM^{D}_{BpsB}$ | pET-24a(+) backbone with T6-Δ$COM^{D}_{BpsB}$ inserted in MCS with restriction sites *Nde*I/ *Bam*H I | This study |
| pET-24a(+)-$C3_{BpsB}$ | pET-24a(+) backbone with $C3_{BpsB}$ inserted in MCS with restriction sites *Nde*I/ *Bam*H I | This study |
| pET-24a(+)-$C6_{BpsC}$ | pET-24a(+) backbone with $C6_{BpsC}$ inserted in MCS with restriction sites *Nde*I/ *Bam*H I | This study |
| *EryR (Erythromycin resistance) *KanR (Kanamycin resistance) | | |

**Table S4|Strains used in this study.**

| Strain | Description | Source |
|---|---|---|
| *A. balhimycina DSM 44591* | wildtype strain | Nadkarni et al.[2]; Wink et al.[3] |
| *A. balhimycina DB1* | Deletion of C-terminal $COM^{D}_{BpsA}$ encoding sequence of the NRPS gene *bpsA* from the balhimycin biosynthetic gene cluster | This study |
| *A. balhimycina DB2* | Deletion of C-terminal $COM^{D}_{BpsB}$ encoding sequence of the NRPS gene *bpsB* from the balhimycin biosynthetic gene cluster | This study |
| *A.balhimycina DB1-2* | Deletion of C-terminal $COM^{D}_{BpsA}$ and $COM^{D}_{BpsB}$ encoding regions of the NRPS gene *bpsA* and *bpsB* from the balhimycin biosynthetic gene cluster | This study |
| *E. coli* JM110 | *lacY dam dcm* [F' *lacIq* Z Δ M15] (Methylation deficient strain ) | Yanisch-Perron et al.[4] |
| *E. coli* ET12567 | F− *dam-13*::Tn*9 dcm-6 hsdM hsdR* (Methylation deficient strain) | MacNeil et al.[5] |
| *E. coli* NovaBlue | Cloning host for plasmid generation: *endA1 hsdR17* (rK12– mK12+) *supE44 thi-1 recA1 gyrA96 relA1 lac F'[proA+B+ lacIqZΔM15*::Tn*10*] (TetR) | Novagen® |
| *E. coli* BL21 (DE3) | Target strain for protein expression | New England Biolabs® |
| **TetR (Tetrycyclin resistance) *StrR (Streptomycin resistance)* | | |
